# Large- and Small-Animal Studies of Safety, Pharmacokinetics (PK), and Biodistribution of Inflammasome-Targeting Nanoligomer in the Brain and Other Target Organs

**DOI:** 10.1101/2024.02.07.579323

**Authors:** Sydney Risen, Breonna Kusick, Sadhana Sharma, Vincenzo S. Gilberto, Stephen Brindley, Mikayla Aguilar, Jared M. Brown, Stephanie McGrath, Anushree Chatterjee, Julie A. Moreno, Prashant Nagpal

## Abstract

Immune malfunction or misrecognition of healthy cells and tissue, termed autoimmune disease, is implicated in more than 80 disease conditions and multiple other secondary pathologies. While pan-immunosuppressive therapies like steroids offer some relief for systemic inflammation for some organs, many patients never achieve remission and such drugs do not cross the blood-brain barrier making them ineffective for tackling neuroinflammation. Especially in the brain, unintended activation of microglia and astrocytes is hypothesized to be directly or indirectly responsible for Multiple Sclerosis (MS), Amyotrophic Lateral Sclerosis (ALS), Parkinson’s Disease (PD), and Alzheimer’s Disease (AD). Recent studies have also shown that targeting inflammasome and specific immune targets can be beneficial for these diseases. Further, our previous studies have shown targeting NF-κB and NLRP3 through brain penetrant Nanoligomer cocktail SB_NI_112 (abbreviated to NI112) can be therapeutic for several neurodegenerative diseases. Here we show safety-toxicity studies, followed by pharmacokinetics (PK) and biodistribution in small- (mice) and large-animal (dog) studies of this inflammasome-targeting Nanoligomer cocktail NI 112. We conducted studies using four different routes of administration: intravenous (IV), subcutaneous (SQ), intraperitoneal (IP), and intranasal (IN), and identified the drug concentration over time using inductively coupled plasma mass spectrometry (ICP-MS) in the blood serum, the brain (including different brain regions), and other target organs like liver, kidney, and colon. Our results indicate the Nanoligomer cocktail has a strong safety profile, and shows high biodistribution (F ∼0.98) and delivery across multiple routes of administration. Further analysis showed high brain bioavailability with a ratio of NI112 in brain tissue to blood serum ∼30%. Our model accurately shows dose scaling, translation between different routes of administration, and interspecies scaling. These results provide an excellent platform for human clinical translation and predicting therapeutic dosage between different routes of administration.

---

The human immune system is the key line of defense in the body against infections and disease and is critical for our survival. Misrecognition of healthy cells, tissue, and organs by the immune system causes more than 80 chronic conditions and these autoimmune diseases affect more than 24 million people in the US alone, with rising prevalence.^1,2^ These diseases disproportionality affect women and are among the leading causes of death amongst young- and middle-aged women.^1,3^ Besides being directly implicated in neurological diseases related to motor function like Multiple Sclerosis (MS) and Amyotrophic Lateral Sclerosis (ALS), several other neurodegenerative diseases like Parkinson’s Disease (PD) and Alzheimer’s Disease (AD) are now hypothesized to be linked to non-specific microglial activation,^4–7^ the resident immune cells in the brain. While there are no available cures for these autoimmune diseases and more specifically for neuroinflammation, pan-immunosuppressive steroids, and other non-specific therapies are utilized for managing symptoms and prolonging disease progression or symptom manifestation for systemic diseases. However, such non-specific immune suppressive methods have significant side effects, especially systemic effects while treating long-term chronic conditions.^8^ Therefore, more targeted immune modulation^9,10^ as well as detailed pharmacokinetic and route of administration studies are necessary for corticosteroids and pan-immunosuppressive therapies to manage adverse effects and contraindications.^8^

## Specific Immune Targeting can Minimize Side-Effects

Selective inhibition of key inflammatory nodes/targets such as TNF, IL-17, IL-23, and IL-6 have improved the remission rates and replaced standard-of-care for a growing number of autoimmune diseases.^9,10^ While recent advances in even more target-specific therapies such as tyrosine kinase 2 (TYK2) instead of Janus kinase (JAK) inhibitors offer further hope in reducing side-effects (e.g. abnormal changes in blood cell, cholesterol, triglyceride levels, etc.),^11^ use of more traditional pharmaceutical modalities such as small-molecule-based targeting is slow and low-throughput with mixed safety profile. On the other hand, high-throughput approaches such as design-based antisense oligonucleotides (ASOs)^12^ can potentially improve translation rates, but suffer from delivery challenges, especially for neurological diseases,^13,14^ and accumulation in first-pass organs.^15,16^ The emergence of new RNA-targeting antisense modalities such as Nanoligomers can have the combined advantage of high delivery and biodistribution profile of small molecules, extremely high target-binding affinity (K_D_ ∼3-5 nM) like antibodies, with the specificity and ease of design using nucleic acid-targeting.^17–23^ Nanoligomers are peptide nucleic-acid-based RNA therapeutic modality that have shown targeted up-and down-regulation of proteins, with extremely high specificity, minimal off-targets, and a high safety profile.^17–23^ However, more detailed safety-toxicity and biodistribution studies need to be conducted across small- and large-animal models for these new modalities to gain greater acceptance towards human clinical translation and pharma adoption.

## Small- and Large-Animal Studies for Safety and Biodistribution and Intraspecies Scaling

While rodents are acceptable small-animal models for autoimmune and neurological disease modeling and induction, dogs are the preferred large-animal model for clinical drug translation to further assess the safety profile, central nervous system (CNS) pharmacokinetic (PK)/pharmacodynamic (PD), and biodistribution parameters.^24^ We conducted extensive small- and large-animal studies for our targeted inflammasome inhibitor, an NF-κB and NLRP3 inhibitor combination (herein described as SB_NI_112 or NI112 in short) as the active pharmaceutical ingredient (API),^17–22^ to assess safety-toxicity, develop a PK and biodistribution model to obtain relevant pharmacological parameters, and obtain insights for interspecies scaling for further translation. We observed extremely high biodistribution parameters across different routes of administration (biodistribution parameter F >0.98), excellent drug delivery across the blood-brain barrier (BBB) (brain tissue concentration ≥ 25-30% of blood serum bound concentration), rapid absorption across different brain sections, lack of accumulation in first-pass organs, as well as renal clearance of unbound/excess drug (or missense molecules).^19,20^ These pharmacological results prove the potential for further translation and the need for the adoption of new therapeutic modalities for the benefit of millions of patients across multiple autoimmune and CNS diseases.

## RESULTS AND DISCUSSION

We utilized four different routes of administration: intravenous (IV), subcutaneous (SQ), intraperitoneal (IP), and intranasal (IN) for a comprehensive assessment of safety-toxicity and biodistribution profile in both small-animal (C57BL/6 mice) and large-animal (beagle dog) models. We explored dose-scaling in different routes, biodistribution parameters, and interspecies scaling, reported in this work.

### Safety-Toxicity Assessment: Long-Term Repeat Dosing, 14-day Acute (5x MTD) Dosing in Small- and Large-Animals for CNS

We first started with a maximum tolerable dose (MTD) assessment of the NF-κB and NLRP3 inhibitor combination NI112 (*nfkb1* (Sequence: AGTGGTACCGTCTGCTA) and *nlrp3* (Sequence: CTTCTACTGCTCACAGG)). Prior studies have shown assessment of NI112 in relevant disease models (MS, AD, Prion/Creutzfeldt-Jacob Disease, neuroinflammation, and autoimmune diseases),^17,18,22,23,25,26^ and Nanoligomer selectivity and high-binding affinity (K_D_ 3.17 nM,^19,20^ other PNA/RNA and PNA/DNA binding studies showed K_D_ 5-8 nM^27^) along with rapid biodistribution and clearance. We observed strong safety-toxicity profile in both animal models for longer-term repeat dosing (IP, 150 mg/kg, 3x a week for 12 weeks, **Fig. 1A**, **B, S1, S2**) as well as single-dose acute toxicity (5x maximum tolerable dose or MTD for IV) followed by 14-day toxicity assessment and clinical monitoring (**Fig. S3, 1C-F, S4, S5)**. An important finding was that a single route (IV) showed a measurable MTD (100 mg/kg in mice, between 15-65 mg/kg in dogs), while other routes showed values well over 450-500 mg/kg, beyond the limit of testing parameters due to solubility limits. As shown in MTD **Table I** below and **Fig. S3,** the use of missense Nanoligomers (dog NI112 in mice) also showed no tolerability issues for 450-500 mg/kg IV. IV route demonstrates extremely rapid delivery and uptake of Nanoligomers in the brain and other organs (∼ minutes), and some test subjects showed some adverse events (AEs) at high concentrations (75 mg/kg in dogs, **Fig. S3**). However, the PK and biodistribution of other routes are slightly slower, leading to better tolerance for high drug-dosing. This along with lack of such MTD for missense administration suggests that the AEs are related to the speed/rapid uptake and target inhibition in the brain and other organs, rather than potential side-effects of the API/drug. The lack of such threshold for missense Nanoligomer also suggests the specificity of target engagement by the drug. We note during dog studies, one subject animal showed acute injection site pain, swelling, and discomfort at 75 mg/kg SQ dosing, which was traced back to the high pH of SQ injection^28^ due to the use of unbuffered saline media to dissolve dried NI112 powder. However, all subsequent dosing used phosphate-buffered saline (PBS), and no animal showed any issues with site injection pain or swelling.

**Fig. 1.**
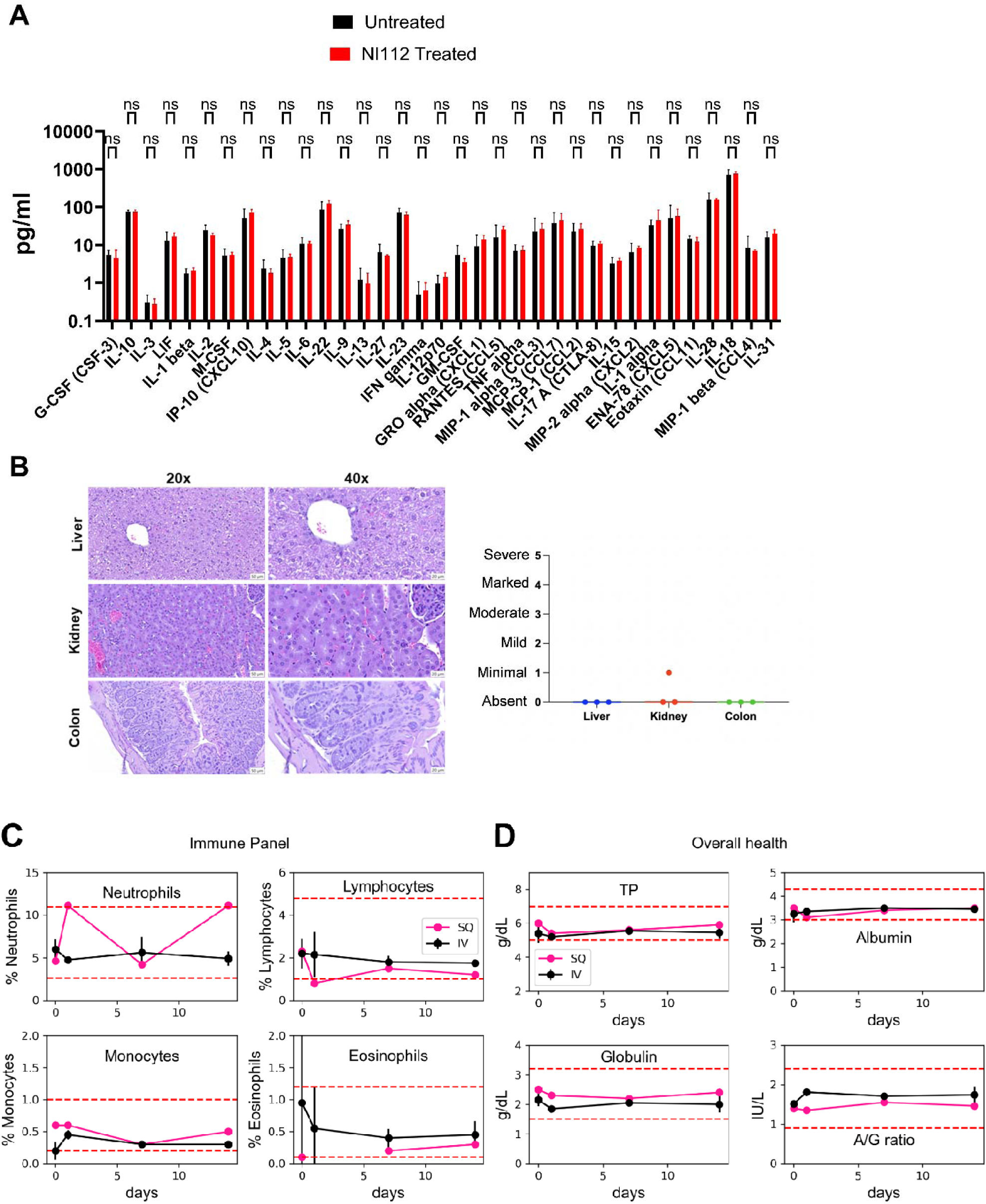
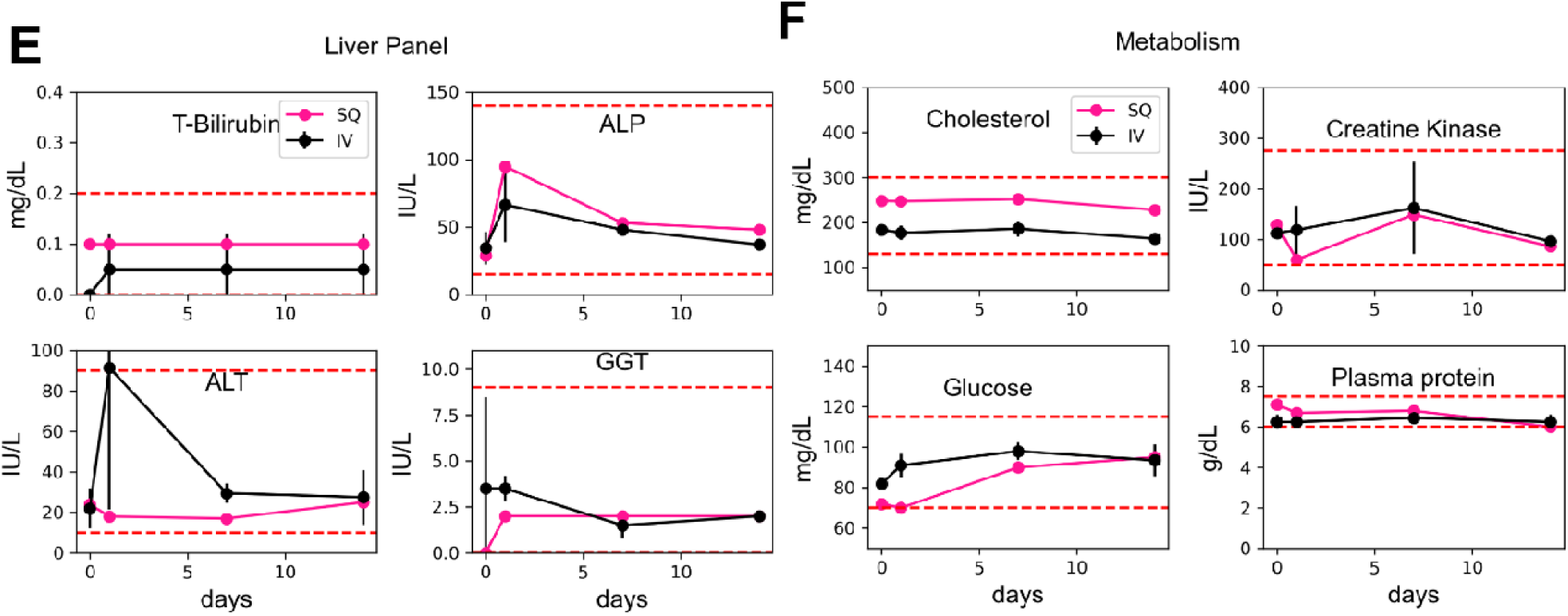
Long-term safety-toxicity and acute single-dose studies reveal the high safety profile of NI112 Nanoligomer. **A)** 12-week dosing of NI112 3x a week shows no change in the immune profile of mice blood serum, as shown using 36-plex cytokine and chemokine markers, measured for treated and untreated animals (n=4/group), ns = non-significant using unpaired t-test for each cytokine. **B)** H&E-stained histology of kidney, liver, and colon samples from mice after high-dose (150 mg/kg 3x a week) long-term (12-weeks) treatment shows no immune cell infiltration or any immunogenic response. Histologic findings in each tissue were diagnosed and graded for severity on a 0-5 scale based on inflammation and hemorrhage; 0=absent, 1=minimal (10% of tissue affected), 2=mild (10-25% of tissue affected), 3=moderate (26-50% of tissue affected), 4=marked (51-75% of tissue affected), 5=severe (75% of tissue affected). There were no significant histologic findings/differences in the liver, kidney, or colons of mice treated with NI112. Kidney nephrons displayed no evidence of inflammation within the glomeruli, supporting the renal clearance shown previously for Nanoligomer molecules.^19,20^ **C)** Immune panel studies of dogs treated with 75 mg/kg NI112 Dog which corresponds to 500 mg/kg dosing in mice, and is well above the MTD. Dogs were monitored for 2 weeks post-administration. The results show a lack of any immunogenic response, and no change in neutrophils, lymphocytes, monocytes, or eosinophils in blood. **D)** Similar assessment of overall health using total protein (TP), albumin (A), globulin (G), and A/G ratio. **E)** Assessment of the liver panel of these animals, and **F)** Metabolism. The red dashed lines show upper and lower normal limits for each measurement. The curves show mean values and bars show standard deviation (Mean ± Standard Deviation, SD). These extensive studies show minimal side effects and a lack of long-term health concerns even at such high acute doses.

**Fig. 2.**
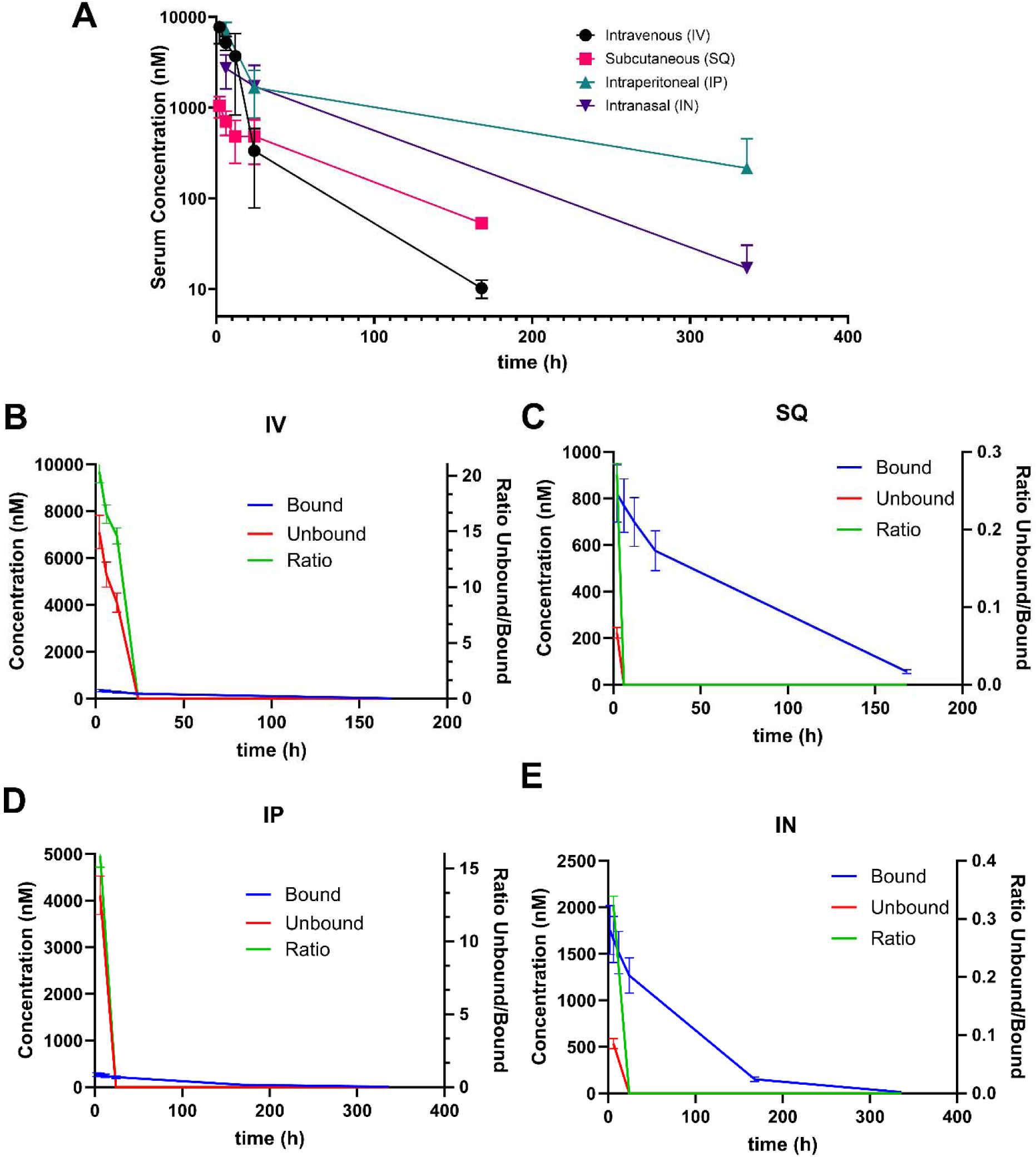
Bound and Unbound API NI112 with time using different routes of administration. **A)** Mice studies showing NI112 concentration in blood serum with time using 4 different routes of administration at MTD IV (100 mg/kg), IP (500 mg/kg), SQ (450 mg/kg), and IN (500 mg/kg). The multiexponential decay with time was used to derive bound and unbound API (NI112) concentration with time using **B)** IV; **C)** SQ; **D)** IP; and **E)** IN route. Right Y axis shows the ratio of unbound/bound API in blood serum. The curves show Mean values, with the bars showing Standard Deviation (Mean ± SD). Number of animals n=4/group for IV and SQ, n=6/group for IN and IP route of administration.

**Table I:**
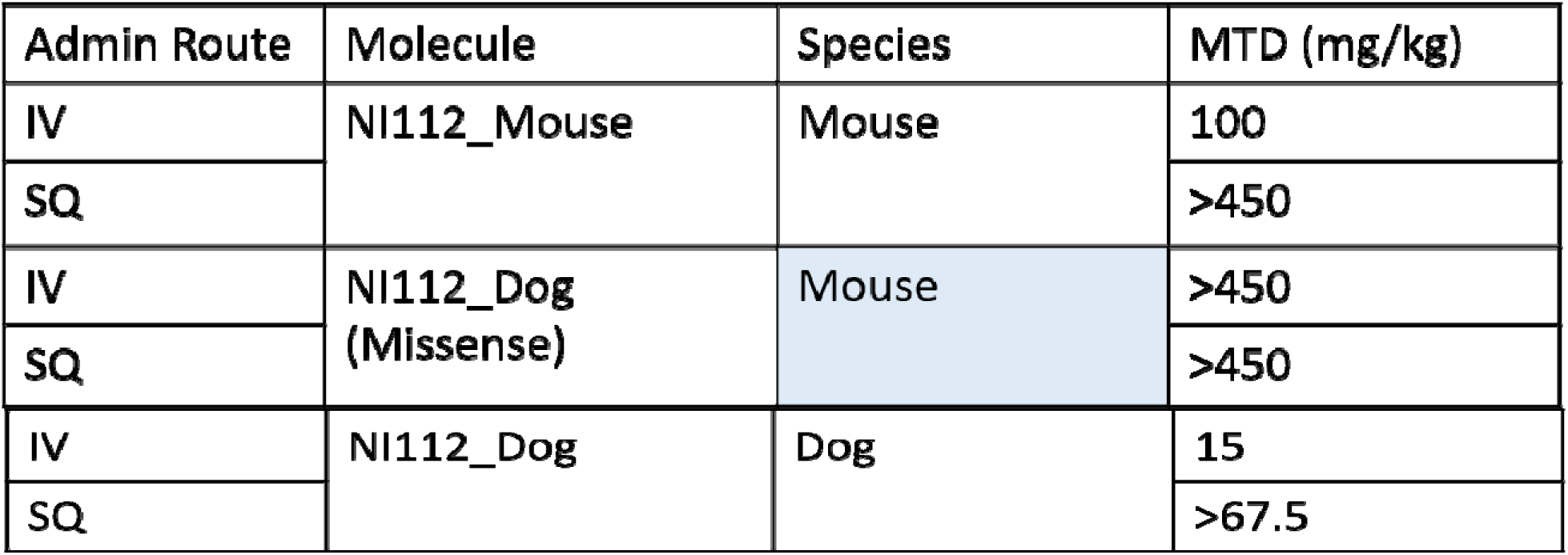
MTD measurements for Mice and Dog Studies Using IV and SQ route of Administration. n=4 for mice studies, n=1-3 for dog studies. 1-test subject each in mice and dog studies showed side-effect/tolerability limit with the IV route, the mouse had experienced seizure-like behavior and the dog exhibited excess salivation and vomiting.

| Admin Route | Molecule | Species | MTD (mg/kg) |
| --- | --- | --- | --- |
| IV | NI112_Mouse | Mouse | 100 |
| SQ |  |  | >450 |
| IV | NI112_Dog (Missense) | Mouse | >450 |
| SQ |  |  | >450 |
| IV | NI112_Dog | Dog | 15 |
| SQ |  |  | >67.5 |

Using these MTD values, we conducted two types of repeat-dosing (using IP) and acute-dosing tests (using IV and SQ). Due to rapid aging in mice, we used 13–15-week-old mice for longer-term (12-week) repeat dosing (3x a week, for chronic diseases) with IP injections, to assess any biochemical or histological changes compared to sham-treated control animals. As shown in **Fig. 1A**, we conducted a detailed blood serum evaluation using 36-cytokine and chemokine biomarkers in blood, for any potential immunological or adverse outcomes (through clinical/health monitoring). We did not observe any statistical change (detailed p-values shown in **Fig. S1**) in any biomarker. Further, we conducted a detailed histological evaluation and scoring for the first pass and clearance organs (liver, kidney, colon, **Fig. 1B, S2**) for chronic-dosed animals, essential for drug metabolism and excretion. We did not observe any evidence of accumulation, organ damage, toxicity, or any immune cell infiltration, as shown using H&E staining. Also, in long-term studies, there was no evidence of accumulation or resulting inflammation in the medulla or nephrons (within glomeruli, **Fig. 1B**), supporting renal clearance for Nanoligomer molecules.^19,20^ These results again demonstrate the high safety profile of the drug for repeat dosing over longer periods.

Next, we conducted acute dose toxicity monitoring in large-animal models (dogs), using 75 mg/kg dose IV administration (5x MTD for IV route). Using the body surface area (BSA) scaling factor canine equivalent dose (CED) (mg/kg) = Mouse dose (mg/kg) * (Mouse Km/Canine Km)) where Km is the BSA scaling factor, to estimate the canine equivalent dose from mouse studies.^29^ This dose factor for safety-toxicity and PK analysis was 0.15.^29^ We assessed the impact of extremely high doses (e.g. 5x MTD using IV administration) or potential acute drug toxicity in a large-animal model (dogs). We monitored these animals over the recommended 14-day period, with extensive clinical sign monitoring and health checkups (**Fig. S3**), and detailed biomarker assessment, as shown in **Fig.1C-F**. We dosed dogs using IV and SQ at 75 mg/kg dose. While few test subjects showed some short-term AEs noted in **Fig. S3**, and 75 mg/kg dose was marked as above MTD in our studies, we sought to further evaluate any adverse health impact. Using a battery of complete blood count (CBC)/chemistry tests for potential immunogenic response (by monitoring neutrophils, lymphocytes, monocytes, or eosinophils in blood, **Fig. 1C**), overall health (total protein TP, albumin A, globulin G, and A/G ratio, **Fig. 1D**), liver enzymes (T-bilirubin, alkaline phosphatase (ALP), alanine transaminase (ALT), gamma-glutamyl transferase (GGT), **Fig. 1E**), and metabolic markers (like cholesterol, creatine kinase, glucose, and plasma protein, **Fig. 1F**), we did not observe any pathogenic change in any test subject. We also monitored red blood cells, electrolytes (**Fig. S4**), and kidney function (**Fig. S5**) for the acute-dosed beagle dogs to assess any adverse health impact through these biochemical markers. We did not observe any lasting health effects throughout the biomarker assays and the 14-day clinical monitoring. We also repeated these studies at the estimated MTD values for different routes (**Figs. S6-S9**), but again did not observe any abnormal values or clinical concern in any test subject. This showed a well-tolerated, non-toxic, and non-pathogenic safety profile of API Nanoligomer NI112 treatment using extremely high (acute) dosing in large-animals.

### Modeling Biodistribution of the API in Blood Serum and Target Organs

We measured API NI112 concentration (using ICP-MS, see Methods) in blood for 4-different administration routes (IV, IP, IN, and SQ) in mice, with time, and used the multiexponential decay to estimate the time-dependent bound and unbound API concentration (**Fig. 2A**). Briefly, the longtime-decay exponential of blood serum API (bound serum) was assessed as a fraction of the unbound API (**Fig. 2B-E**), and observed the unbound API decays with time, almost to zero, within 24-hours of drug administration in mice. Another important observation was that while IV and IP routes exhibited an appreciable unbound/bound API ratio (**Fig. 2B, D**), SQ and IN routes had a very low ratio of unbound API measured in the blood (**Fig. 2C, E**). Eventually (∼24 hours post-administration) all the measured API is bound. These results are consistent with our prior studies, which tracked any unbound API (and all missense sequences) cleared through urine by the animals.^19,20^ We also used PK solver^30^ to derive pharmacokinetic parameters for the API in blood serum and measurement of biodistribution coefficient (F=AUC _Test route_/AUC _I.V.,_ AUC: Area under the curve), as shown in **Figs. S10, S11**. While IV and SQ routes showed rapid uptake in the blood (T_max_ 2 hours) compared to IP and IN, the target organ brain tissue concentration showed higher variance. The half-life in blood (∼1-2 days) was also significantly lower than in the brain (∼ 1-2 weeks), showing the rapid uptake of API in different organs and drug clearance from the bloodstream through urine.^19,20^

Further measurements of API NI112 were done in the target organ (the brain, **Figs. 3, 4**) and other potential target and/or first pass, and clearance organs (liver, kidney, colon, **Fig. 5**). We found high drug bioavailability (∼30% of the drug) in the brain across the BBB, compared to API concentration in the blood, and much higher values in liver, kidney, and colon. We even observed excellent biodistribution among different brain regions (hippocampus, thalamus, cortex, cerebellum, and brainstem, **Fig. 3C**), providing good confidence for drug delivery and using other Nanoligomer target sequences for different neuronal disease applications. We also developed a model for predicting target organ concentration and precise dosing concentration and dose frequency (**Fig. 4**). The predictive nature of such PK/PD biodistribution profile, combined with detailed disease model studies in small-animals could provide an excellent platform using dose scaling and similar interspecies translation parameters for large-animal model.

**Fig. 3.**
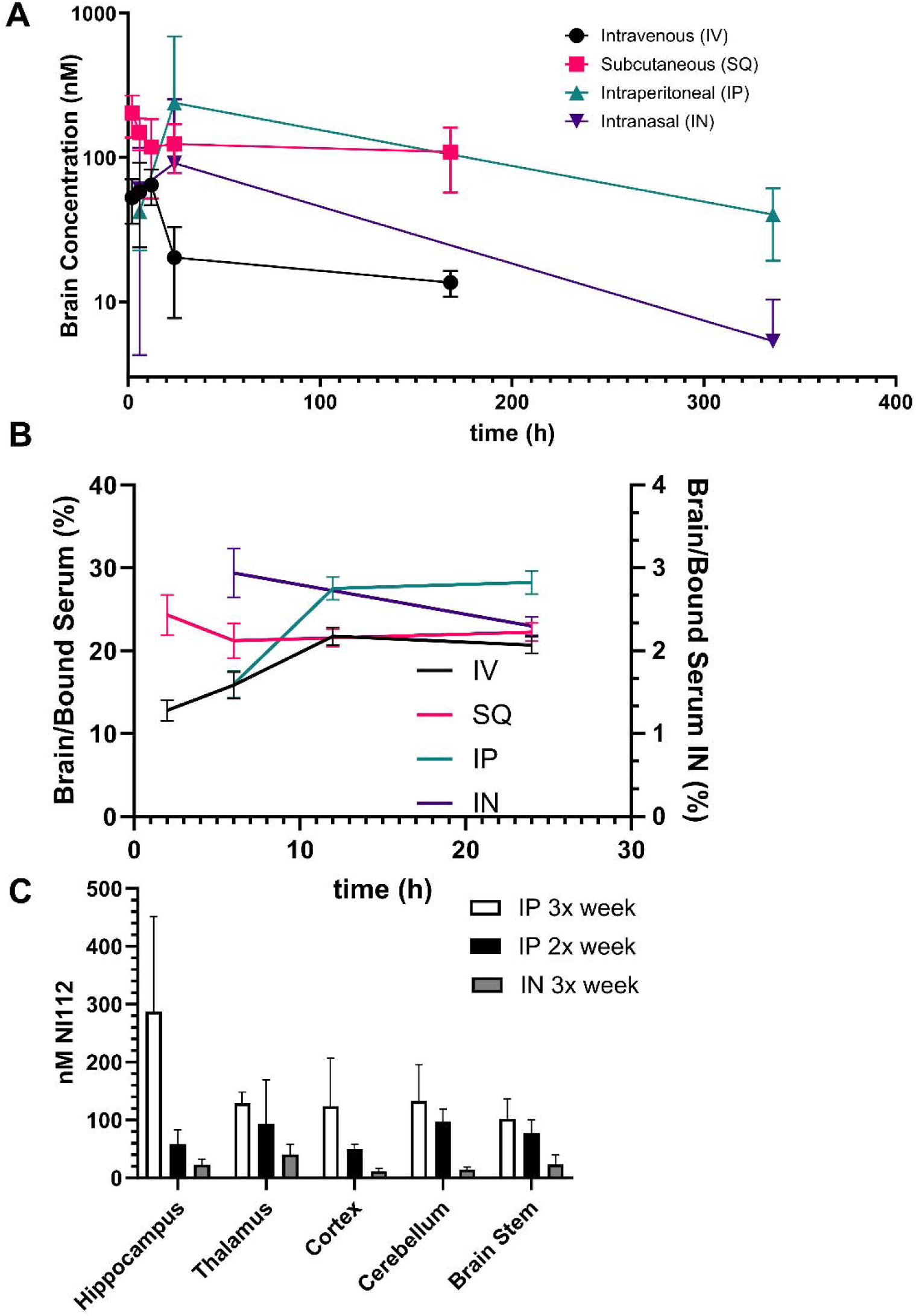
High drug bioavailability and BBB-crossover for neurological disease application. **A)** Mice studies showing NI112 concentration in brain tissue with time using 4 different routes of administration at MTD IV (100 mg/kg), IP (500 mg/kg), SQ (450 mg/kg), and IN (500 mg/kg). **B)** Ratio of Brain tissue to bound API concentration in blood serum shows a fairly consistent value (∼25-30%) for 3 routes of administration IV, IP, and SQ. Right Y axis shows the ratio of brain tissue/bound API in blood serum for IN administration which is one order of magnitude lower (∼3%). **C)** High bioavailability of API (NI112) in different brain regions with different dosing regimens (3x a week and 2x a week) using IP and IN route at 150 mg/kg dose. The curves show Mean values, with the bars showing Standard Deviation (Mean ± SD). Number of animals n=4/group for IV and SQ, n=6/group for IN and IP route of administration.

**Fig. 4.**
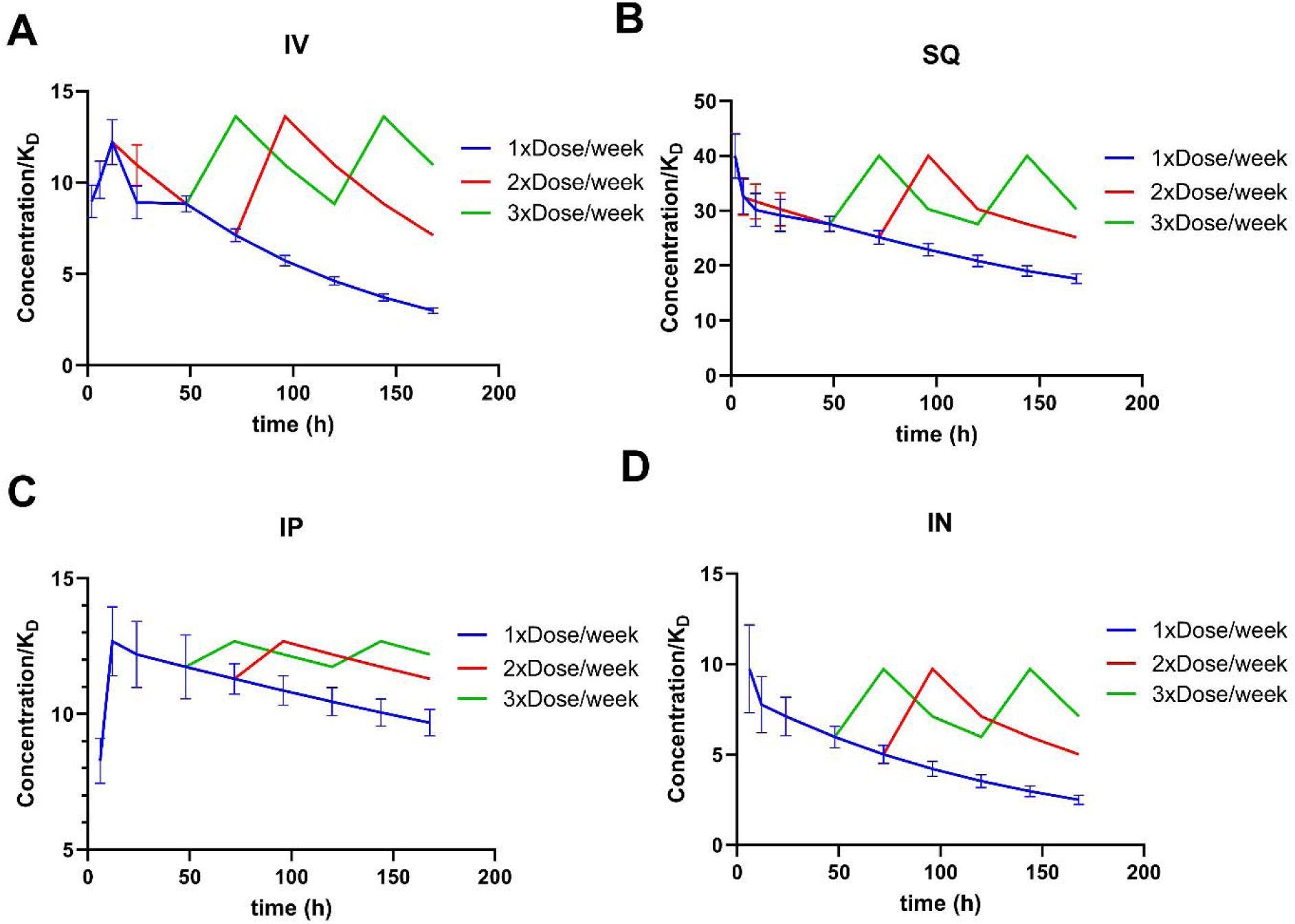
Dosing and repeat dose modeling data for small-animal studies in the target organ (the brain). **A)** IP administration of API NI112 concentration in brain tissue normalized with drug K_D_ (5 nM, Nanoligomer measured K_D_ of 3.37 nM,^19,20^ other PNA studies show values 5-8 nM^27^) with time using IV (100 mg/kg dosing). The blue curve with standard deviation shows the measured values after a single dose at MTD, whereas red (2x a week) and green (3x a week) are modeled values, shown without SE for clarity. **B)** SQ dosing (450 mg/kg) showing single and repeat dosing effects on target organ (the brain) concentration normalized by K_D_. **C)** IP (500 mg/kg) and **D)** IN (500 mg/kg) show high drug bioavailability of API (NI112) in brain regions with different dosing regimens (3x a week and 2x a week). The curves show Mean values, with the bars showing Standard Deviation (Mean ± SD). Number of animals n=4/group for IV and SQ, n=6/group for IN and IP route of administration.

**Fig. 5.**
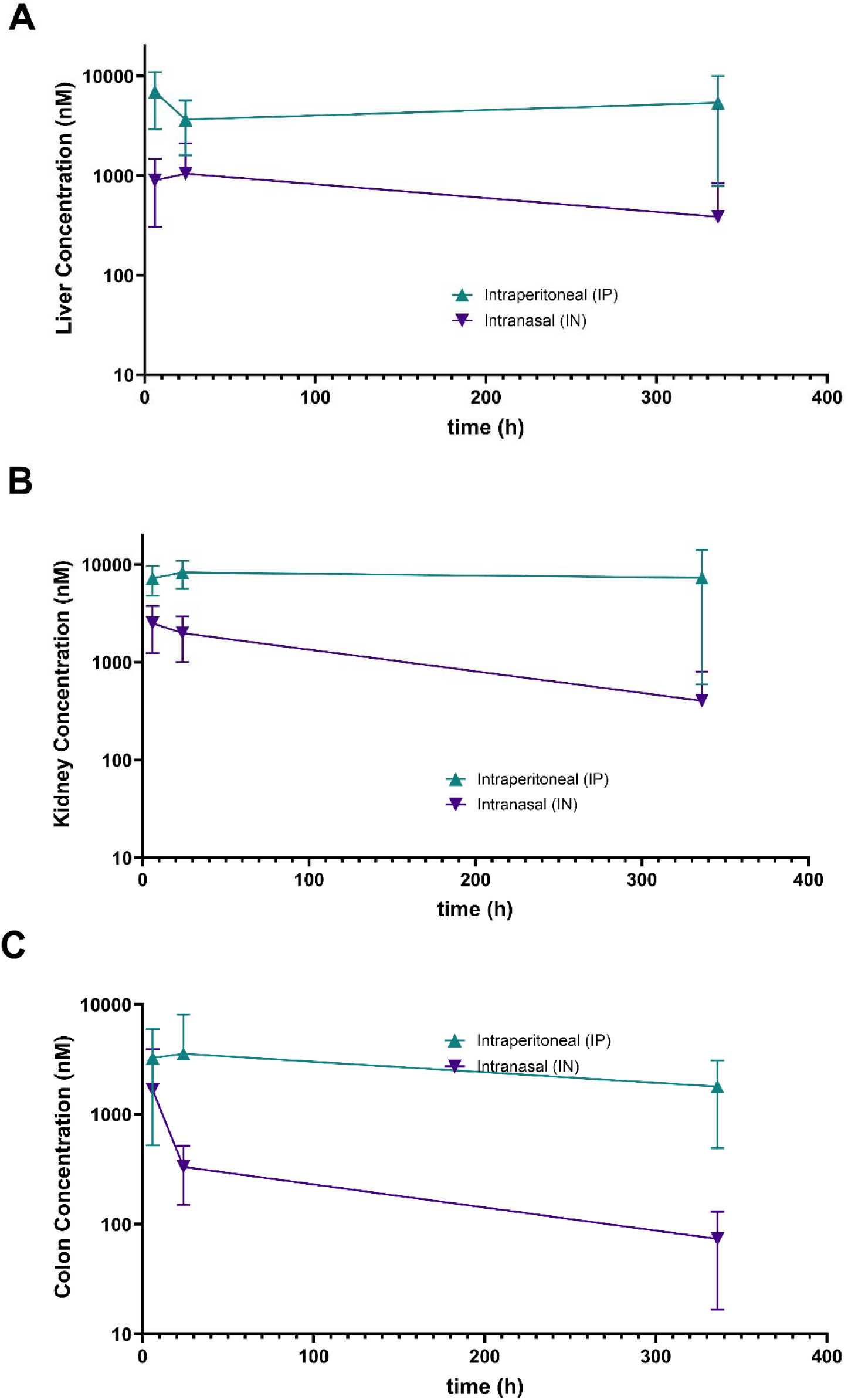
NI112 tissue concentration in the first pass and clearance organs. Mice studies showing NI112 concentration in different first pass and clearance organ tissues with time using IP and IN routes of administration **A)** Liver, **B)** Kidney, and **C)** Colon. The curves show Mean values, with the bars showing Standard Deviation (Mean ± SD). Number of animals n=6/group for IN and IP route of administration.

### Biodistribution in the Large-Animal Model (Dogs)

We measured NI112 concentration with time in blood serum for IV, and SQ dosing in dogs (**Figs. 6-8**). We obtained PK parameters for different routes and concentrations used (**Figs. S10, S12**). The PK parameters showed similar peak API concentration as rodents, but significantly higher bound API concentrations with time, and much higher AUC values, due to higher target mRNA and protein concentration (NF-κB and NLRP3) in dogs than mice, well beyond BSA scaling used. These AUC ratios also correspond with interspecies scaling and are better predictors than BSA estimation (Km ratios) used for dose scaling between mice and dog studies. We also modeled multiexponential decay to estimate the bound and unbound API concentration with different IV (**Fig. 6**) and SQ (**Fig. 7**) doses. Our model showed a strong predictive, linear relationship (ratios/slope slightly lower than 1) between the bound serum API concentration with time and dose ratios (**Fig. 8**). Therefore, our PK parameters and the model showed an excellent biodistribution profile for the API NI112, and very predictive dose scaling for both small- and large-animal models. Therefore, we proceeded to combine our model results (small- and large-animal) to develop interspecies scaling and translation parameters for the API NI112.

**Fig. 6.**
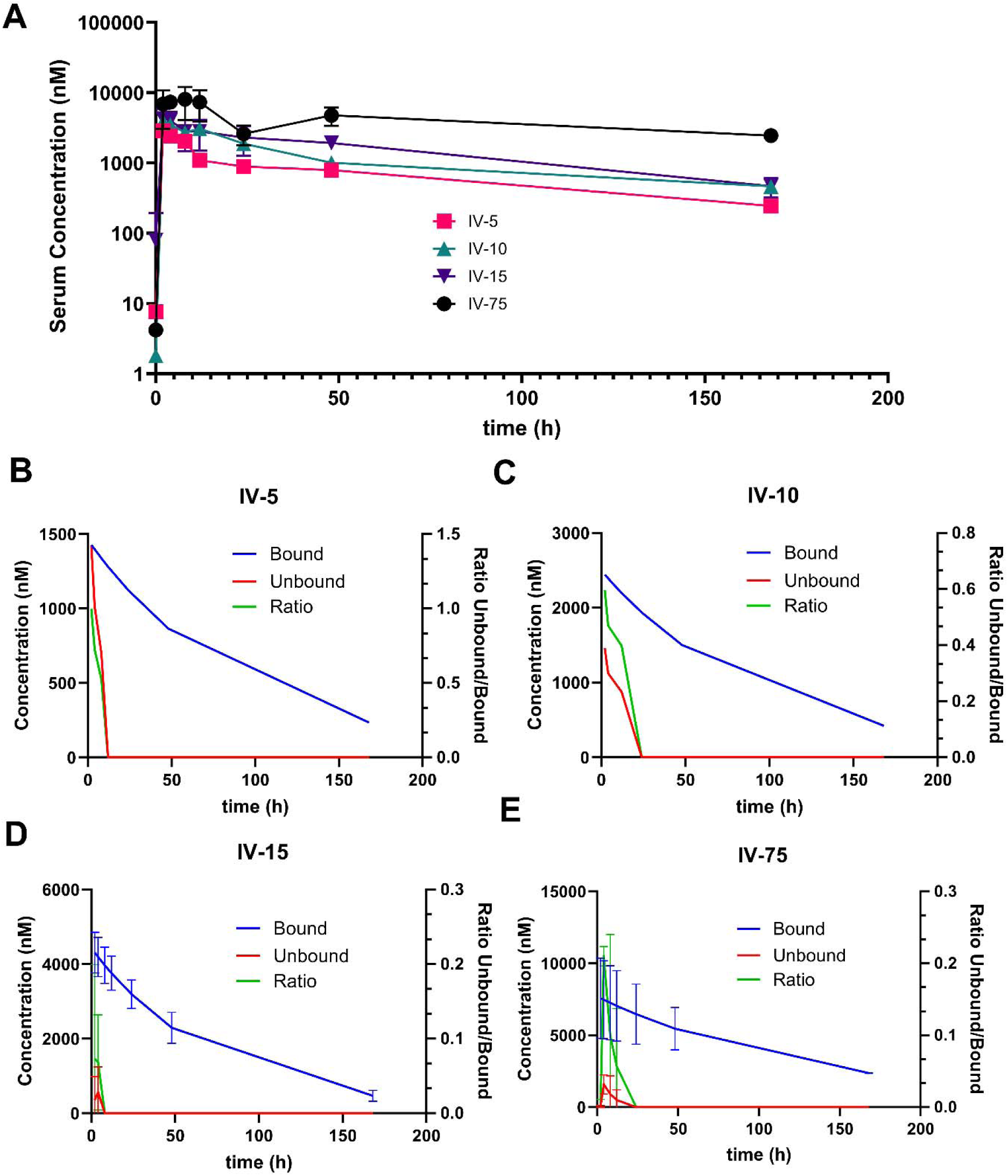
Large-animal dog studies using IV administration of API NI112 at different concentrations. **A)** Blood serum concentration of the API with 5, 10, 15, and 75 mg/kg dosing in beagle dogs. Multiexponential decay of API concentration was used to determine bound and unbound serum concentrations for: **B)** 5 mg/kg; **C)** 10 mg/kg; **D)** 15 mg/kg (MTD, n=3/group); and **E)** 75 mg/kg (5xMTD) IV dose. The blue curve (bound) and red curve (unbound) show API in blood, along with the Ratio of unbound/bound API (green curve, shown on Right Y-axis). The curves show Mean values, with the bars showing Standard Deviation (Mean ± SD).

**Fig. 7.**
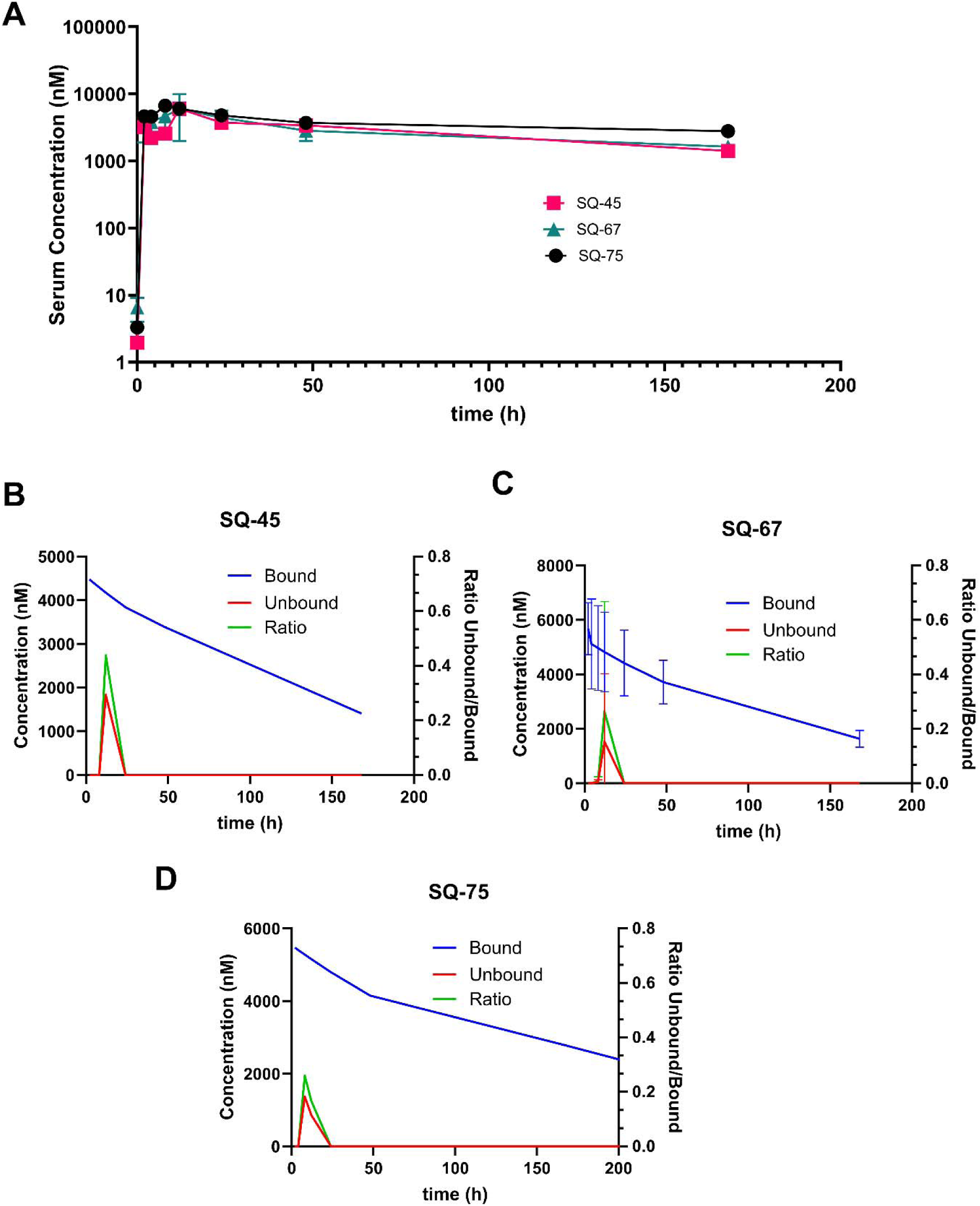
Large-animal dog studies using SQ administration of API NI112 at different concentrations. **A)** Blood serum concentration of the API with 45, 67, and 75 mg/kg dosing in beagle dogs. Multiexponential decay of API concentration was used to determine bound and unbound serum concentrations for: **B)** 45 mg/kg; **C)** 67 mg/kg; **D)** 75 mg/kg SQ dose. The blue curve (bound) and red curve (unbound) show API in blood, along with the Ratio of unbound/bound API (green curve, shown on Right Y-axis). The curves show Mean values, with the bars showing Standard Deviation (Mean ± SD).

**Fig. 8.**
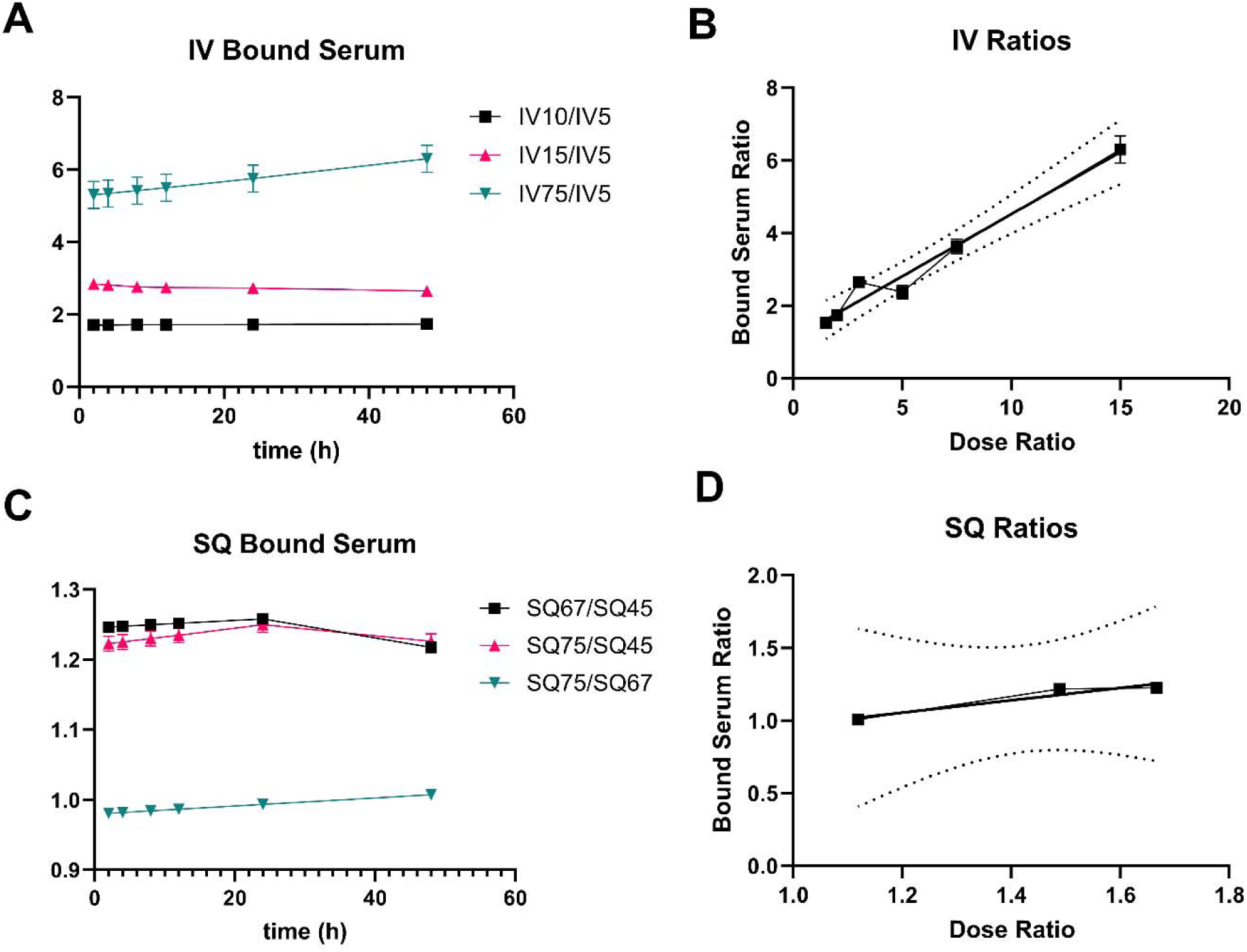
Dose scaling in dog studies using IV and SQ administration of API NI112 at different concentrations. **A)** IV concentration of bound serum shows a nearly constant ratio with the lowest concentration (5 mg/kg). The ratio of 75/5 mg/kg (inverted triangle), 15/5 mg/kg (upward triangle), and 10/5 mg/kg (square) shows flatline values. **B)** Linear regression between the bound serum ratio and the dose ratio, with dashed curves showing the 5% and 95% confidence intervals. **C)** Bound serum ratio between respective SQ doses. **D)** Linear regression between bound serum ratio and dose ratio, showing easily predictable dose scaling that can be used to determine dosing and target organ concentrations in large-animals (beagle dogs). The curves show bound API from Mean values, and the bars show Standard Deviation (Mean ± SD).

### Translation and Scaling across Species (Between Small- and Large-Animal Models)

Using the detailed PK, biodistribution, and scaling models for large- and small-animals, we attempted to develop interspecies scaling and translation parameters. While the proposed BSA scaling utilized the Km ratio as a simplistic model,^29^ several studies have highlighted the inadequateness of such simple model scaling and the need for more concrete (area under the curve AUC) and other PK and biodistribution-based modeling parameters to scale.^31^ We observed that even though the doses were BSA-scaled between large-animal (dog) and small-animal (mice) studies for both IV and SQ administration of API based on Km (Km dose factor for dog/mice was 0.15), the value we observed for two routes was not one (**Fig. 9**). Using the ratio of these scaled doses between species, we observed a clear ratio of 5-6.5 for SQ, and between 12-15 for IV administration, presumably based on accessible target ratio (NF-κB and NLRP3 protein amount between species) and AUC ratios (**Fig. 9**). Our previous studies showed that 150 mg/kg NI112 IP dosing in mice was therapeutic in several neurological diseases.^17,22,23^ Based on dose scaling for different routes in mice shown here (**Fig. 4**), it corresponds to 50 mg/kg SQ and 30 mg/kg IV dosing in mice. For interspecies scaling, using both BSA (factor 0.15) and SQ and IV factors shown in Fig. 9, CED corresponds to 1.5 mg/kg SQ and 0.35 mg/kg IV dosing in dogs, and using BSA scaling from dog to human, an estimated human equivalent dose (HED) of 0.5-0.75 mg/kg SQ and 0.175 mg/kg IV. For a 70 kg human, this roughly translates to therapeutic dosing of 35-52 mg API through SQ and 12.5 mg dosing through IV. We continue to develop a predictive model for further scaling for human translation but can use these models and scaling/translation ratios to provide a better estimate for HED. Together, the developed model and pharmacological biodistribution parameters obtained provide a good foundation for further translation and assessment of the API.

**Fig. 9.**
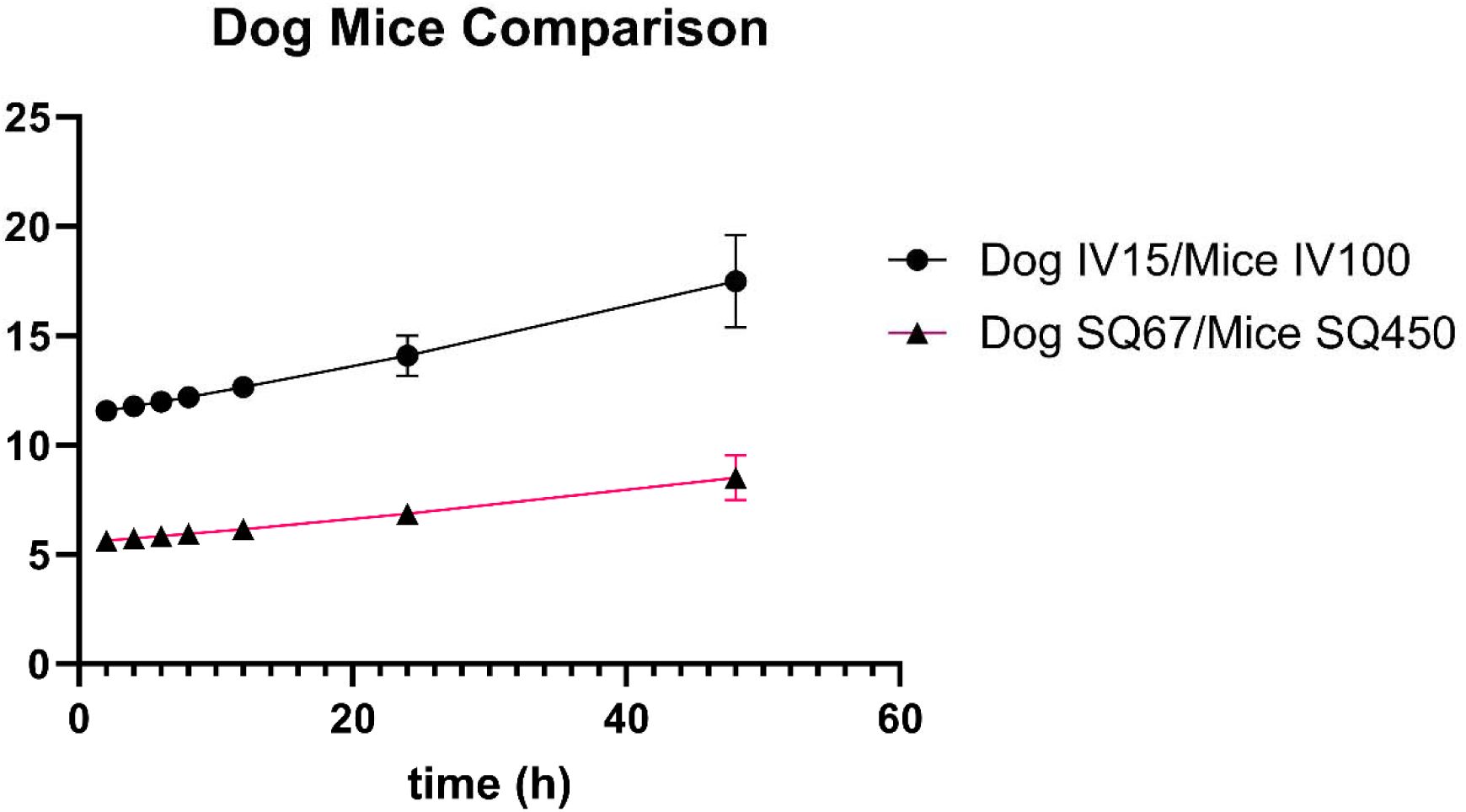
Scaling between large-animal (dog) and small-animal (mice) studies using IV and SQ administration of API NI112 at different concentrations. The doses chosen were based on the Km scaling described above (CED (mg/kg) = Mouse dose (mg/kg) * (Mouse Km/Canine Km)) to estimate the canine equivalent dose from mouse studies. The dose factor for dogs/mice was 0.15.^29^ The curves show bound API from Mean values, and the bars show Standard Deviation (Mean ± SD).

## CONCLUSIONS

In conclusion, we conducted a detailed safety-toxicity assessment (long-term repeat dosing and extremely high 5x MTD acute dosing) in small- (C57BL/6 mice) and large-animal (beagle dogs) models for assessment of neurotherapeutic API NI112. We assessed histological, detailed biochemical (through multiplexed ELISA), and extensive CBC/chemistry biomarker monitoring of immune, metabolic, liver, kidney, and cardiac function. Not a single test subject showed any lasting health impact of the drug administration, and no evidence of any immunogenic response, drug accumulation, tissue damage, or immune cell infiltration was observed. Overall, the animals showed that the new Nanoligomer modality has a strong safety profile. We also conducted detailed PK and biodistribution studies using different routes of administration (IV, SQ, IP, IN) at different doses, and obtained relevant PK parameters. We observed high biodistribution across the target organ the brain (∼30% of API in the brain across BBB, compared to that bound in blood serum), and across the different brain regions (the hippocampus, thalamus, cortex, cerebellum, and the brain stem) and high therapeutic bioavailability (>15-40 K_D_) for low/moderate dosing, and effective target engagement. We found good agreement between administered dose ratios and measured API across the blood serum, the brain, and other first-pass and clearance organs, as well as good agreement between interspecies parameters obtained. While the simple BSA scaling did not provide directly translatable doses (BSA-scaled doses across species do not result in ratios of 1), the values obtained here can provide an excellent starting point/platform for further testing and clinical translation of the API for inflammatory and CNS diseases.

## METHODS

### Nanoligomer Synthesis

NFκB and NLRP3 antisense Nanoligomers were synthesized at Sachi Bio following established methods detailed elsewhere.^17–22^ Briefly, the 17-base-long peptide nucleic acid (PNA)^17,18,21,32^ sequence was selected to optimize solubility and specificity, targeting the start codon regions of *nfkb1* (Sequence: AGTGGTACCGTCTGCTA) and *nlrp3* (Sequence: CTTCTACTGCTCACAGG), within the Mouse genome (GCF_000001635.27), identified using our bioinformatics toolbox.^21^ The PNA was synthesized on a Vantage peptide synthesizer (AAPPTec, LLC) using solid-phase Fmoc chemistry. Fmoc-PNA monomers, wherein A, C, and G monomers were shielded with Bhoc groups, were obtained from PolyOrg Inc. Post-synthesis, the peptides were attached to gold nanoparticles and subsequently purified using size-exclusion filtration. The conjugation process and the concentration of the purified Nanoligomer was measured using absorbance at 260 nm (for PNA detection) and 400 nm (for nanoparticle quantification).

### In vivo mice studies

Adult female and male C57Bl6/J mice purchased from Jackson Laboratories were housed throughout all experiments at ∼18-23°C on a 12-light/12-dark cycle. Fresh water and *ad-libitum* food (Tekland 2918; 18% protein) was routinely provided to all cages. Animals were consistently health checked by the veterinary staff at Colorado State University (CSU). This protocol was approved by the CSU Institutional Animal Care and Use Committee (IACUC) and Laboratory Animal Resources staff.

### Tissue collection

Mice were culled after deep anesthetizing with isoflurane, and ∼1 ml of blood was removed via cardiac puncture followed by cervical dislocation. The brain was sliced sagittally and the right side of the brain was fixed in 10% normal buffered formalin (NBF) and the left hippocampus, thalamus, brainstem, and a piece of the left cortex and cerebellum tissue were removed, flash-frozen, and stored at -80°C until further processing. The liver, kidney, and colon were also removed, fixed in 10% NBF, and processed (paraffin-embedded) for immunostaining (details below).

### Large-animal dog studies

All aspects of the study were approved by the CSU IACUC (protocol ID 4320, approved April 18, 2023). Fifteen healthy, purpose-bred, mixed-sex (14 female, 1 male) beagles between 2 and 3 years of age were enrolled. Dogs weighed a mean of 8kg with a range of 6.7 to 10.6kg. All animals were housed in an on-site research facility and were fed, cleaned, socialized, and examined daily by Laboratory Animal Resource staff or veterinarians. Due to animal temperament and intergroup aggression, dogs were housed in 1 group of 5 dogs, 2 groups of 3 dogs, and 2 groups of 2 dogs. Each dog was identified by a distinct ear tattoo. Dogs were offered water continuously and were fed Exclusive Adult Dog Chicken & Brown Rice Formula dry food in the mornings. Blood samples were collected at time t=0, 2, 4, 8, 12, 24, 48 hours, 7, and 14 days.

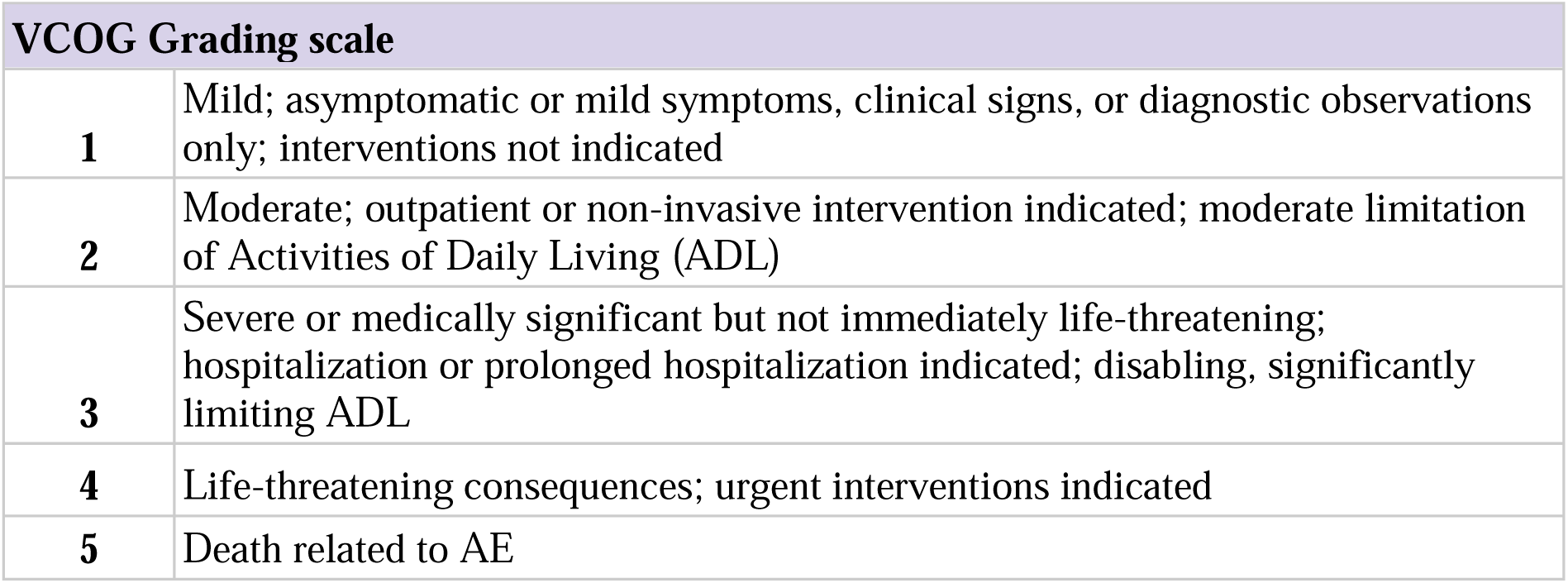

### Multiplexed ELISAs

Blood serum and tissue homogenates from mice were subjected to analysis using the ThermoFisher 36-plex Procartaplex cytokine/chemokine panel. ^17,18^ The process involved homogenizing 25uL of 10 mg/ml brain tissues with ProcartaPlex™ Cell Lysis Buffer (ThermoFisher) and 5-mm stainless steel beads (Qiagen) at 25 Hz for 1.5-3 minutes, utilizing a Qiagen/Retsch Bead Beater. After homogenization, samples underwent centrifugation at 16,000 x g for 10 minutes at 4 °C. Following centrifugation, homogenized samples were evaluated for protein concentration using the DC Protein Assay (Bio-Rad) kit protocol. Protein measurements were acquired through a Tecan GENios microplate reader.

The homogenate was then processed in accordance with the manufacturer’s provided protocol and analyzed using a Luminex MAGPIX xMAP instrument. Standards for each cytokine/chemokine were utilized at 1:4 dilutions (8-fold dilutions), alongside background and controls. Subsequently, sample concentrations were determined from a standard curve using the Five Parameter Logistic (5PL) curve fit/quantification method.

### Inductively coupled plasma mass spectrometry (ICP-MS)

ICP-MS quantification was conducted following established protocols as outlined in prior publications.^19,20^ Briefly, tissue samples underwent homogenization in a TissueLyser II bead mill (Qiagen, Germantown, MD) at 30 Hz for 3 minutes, with the addition of 1 μl deionized water per 1 mg of organ tissue. Homogenates were subsequently digested in 500 μl of aqua regia (3:1 hydrochloric acid to nitric acid) for 4 hours at 100 °C. Pellets resulting from the digestion process were resuspended in water and subjected to analysis using a NexION 2000B single quadrupole ICP-MS system (PerkinElmer, Waltham, MA). The sample introduction utilized a Meinhard nebulizer with a cyclonic glass spray chamber, and a nickel sample and skimmer cone were employed in conjunction with an aluminum hyperskimmer cone. A seven-point linear standard curve was generated within a concentration range of 0-250 ppb Au, and linearity was defined at an R^2^ value of greater than 0.995. The 1000 ppm gold standard solution (Ricca Chemical Company, Arlington, TX), and TraceMetal grade 70% HNO_3_ (ThermoFisher Scientific, Waltham, MA) were used as obtained. All water used was sourced from a Milli-Q system (Millipore, Burlington, MA). Subsequent data analysis was carried out using Microsoft Excel.

### Hematoxylin and Eosin staining for tissue morphology

Brains and visceral organs were fixed in 10% NBF at room temperature for at least 48 hours. Tissues were processed using a Leica TP1020 Automatic Benchtop Tissue Processor and embedded in paraffin wax (Cancer Diagnostics, Cat #: EEPAR56). Tissues were sectioned on a ThermoFisher Scientific HM 325-2 Manual Microtome at 5μm thickness and mounted on positively charged glass slides (Superfrost Plus, Cancer 232 Diagnostics, Cat #: 4951) for staining and analysis. One section per animal was deparaffinized and stained with hematoxylin (Cancer Diagnostics, Cat#: #HTV-4) and eosin (Cancer Diagnostics, Cat#: #ETV) (H&E) for the determination of histopathological changes. Whole tissue images were taken for analysis using an Olympus DP70 camera using an Olympus UPlanSApo 20x objective (N.A. = 0.75).

### Statistics

Statistical tests, numbers of mice, and p-values are indicated in figure legends. GraphPad Prism and Microsoft Excel were used for statistical analysis.

### Ethics Approval

Mice were euthanized by deeply anesthetizing with isoflurane followed by decapitation. All mice were bred and maintained at Lab Animal Resources, accredited by the Association for Assessment and Accreditation of Lab Animal Care International, in accordance with protocols approved by the Institutional Animal Care and Use Committee at Colorado State University. All dog experimentation was also approved by the Clinical Review Board and the Institutional Animal Care and Use Committee at Colorado State University.

## ASSOCIATED CONTENT

### Supporting Information

Supporting figures S1-S12, with p-values for 36-plex Multiplexed ELISA plots for safety-tox evaluation, histology figures for H&E staining in long-term dosing studies, VCOG scoring of dosed animals, CBC-chemistry evaluation and different panels for evaluation of 5xMTD dosing and at MTD dosing, biodistribution analysis, and PK parameters.

### Author Information Corresponding Author

*

### Author Contributions

P.N. conceived the idea, designed the experiments, and synthesized the Nanoligomers. J.A.M. and S.M. designed and supervised animal studies. S.R. and J.A.M. conducted the small-animal mice studies, tissue dissections, immunofluorescence, and scoring. S.S. and V.G. homogenized the tissues, carried out sample digestions and sample preparation for ICP-MS, and conducted ELISA (with P.N.). B.K., M.A., and S.M. conducted all dog studies and sample collections. S.B. and J.B. conducted ICP-MS measurements. P.N. and A.C. wrote the manuscript with input from all the authors. All authors read the manuscript and provided input.

**Notes**

## ACKNOWLEDGMENTS

Authors acknowledge financial support from NASA SBIR Award 80NSSC22CA116.

## TOC IMAGE

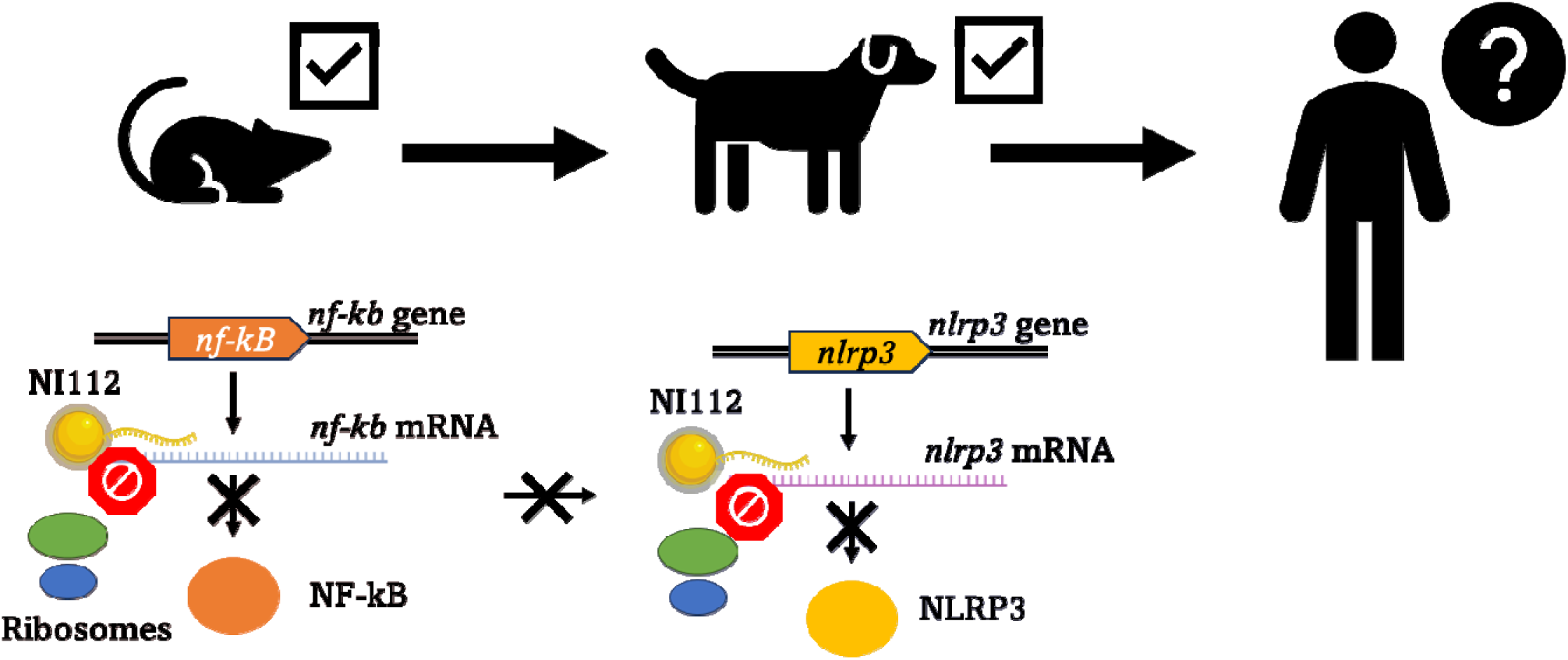

## References

(1) US Department of Health and Human Services. Progress in Autoimmune Diseases Research. Natl. Institutes Heal. 2005, 05–5140 (5).

(2) Cooper, G. S.; Bynum, M. L. K.; Somers, E. C. Recent Insights in the Epidemiology of Autoimmune Diseases: Improved Prevalence Estimates and Understanding of Clustering of Diseases. J. Autoimmun. 2009, 33 (3–4). 10.1016/j.jaut.2009.09.008.

(3) Fairweather, D. L.; Rose, N. R. Women and Autoimmune Diseases. In Emerging Infectious Diseases; 2004; Vol. 10. 10.3201/eid1011.040367.

(4) Heneka, M. T. Inflammasome Activation and Innate Immunity in Alzheimer’s Disease. Brain Pathol. 2017, 27 (2). 10.1111/bpa.12483.

(5) Ising, C.; Venegas, C.; Zhang, S.; Scheiblich, H.; Schmidt, S. V.; Vieira-Saecker, A.; Schwartz, S.; Albasset, S.; McManus, R. M.; Tejera, D.; Griep, A.; Santarelli, F.; Brosseron, F.; Opitz, S.; Stunden, J.; Merten, M.; Kayed, R.; Golenbock, D. T.; Blum, D.; Latz, E.; Buée, L.; Heneka, M. T. NLRP3 Inflammasome Activation Drives Tau Pathology. Nature 2019, 575 (7784). 10.1038/s41586-019-1769-z.

(6) Ní Chasaide, C.; Lynch, M. A. The Role of the Immune System in Driving Neuroinflammation. Brain Neurosci. Adv. 2020, 4. 10.1177/2398212819901082.

(7) Wes, P. D.; Easton, A.; Corradi, J.; Barten, D. M.; Devidze, N.; DeCarr, L. B.; Truong, A.; He, A.; Barrezueta, N. X.; Polson, C.; Bourin, C.; Flynn, M. E.; Keenan, S.; Lidge, R.; Meredith, J.; Natale, J.; Sankaranarayanan, S.; Cadelina, G. W.; Albright, C. F.; Cacace, A. M. Tau Overexpression Impacts a Neuroinflammation Gene Expression Network Perturbed in Alzheimer’s Disease. PLoS One 2014, 9 (8). 10.1371/journal.pone.0106050.

(8) Yasir, M.; Goyal, A.; Sonthalia, S. Corticosteroid Adverse Effects. In StatPearls; 2022.

(9) Jung, S. M.; Kim, W. U. Targeted Immunotherapy for Autoimmune Disease. Immune Network. 2022. 10.4110/in.2022.22.e9.

(10) Villoslada, P.; Moreno, B.; Melero, I.; Pablos, J. L.; Martino, G.; Uccelli, A.; Montalban, X.; Avila, J.; Rivest, S.; Acarin, L.; Appel, S.; Khoury, S. J.; McGeer, P.; Ferrer, I.; Delgado, M.; Obeso, J.; Schwartz, M. Immunotherapy for Neurological Diseases. Clinical Immunology. 2008. 10.1016/j.clim.2008.04.003.

(11) Chimalakonda, A.; Burke, J.; Cheng, L.; Catlett, I.; Tagen, M.; Zhao, Q.; Patel, A.; Shen, J.; Girgis, I. G.; Banerjee, S.; Throup, J. Selectivity Profile of the Tyrosine Kinase 2 Inhibitor Deucravacitinib Compared with Janus Kinase 1/2/3 Inhibitors. Dermatol. Ther. (Heidelb). 2021, 11 (5). 10.1007/s13555-021-00596-8.

(12) Rinaldi, C.; Wood, M. J. A. Antisense Oligonucleotides: The next Frontier for Treatment of Neurological Disorders. Nature Reviews Neurology. 2018. 10.1038/nrneurol.2017.148.

(13) Mazur, C.; Powers, B.; Zasadny, K.; Sullivan, J. M.; Dimant, H.; Kamme, F.; Hesterman, J.; Matson, J.; Oestergaard, M.; Seaman, M.; Holt, R. W.; Qutaish, M.; Polyak, I.; Coelho, R.; Gottumukkala, V.; Gaut, C. M.; Berridge, M.; Albargothy, N. J.; Kelly, L.; Carare, R. O.; Hoppin, J.; Kordasiewicz, H.; Swayze, E. E.; Verma, A. Brain Pharmacology of Intrathecal Antisense Oligonucleotides Revealed through Multimodal Imaging. JCI Insight 2019, 4 (20). 10.1172/jci.insight.129240.

(14) Goto, A.; Yamamoto, S.; Iwasaki, S. Biodistribution and Delivery of Oligonucleotide Therapeutics to the Central Nervous System: Advances, Challenges, and Future Perspectives. Biopharmaceutics and Drug Disposition. 2023. 10.1002/bdd.2338.

(15) Geary, R. S.; Yu, R. Z.; Siwkowski, A.; Levin, A. A. Pharmacokinetic/Pharmacodynamic Properties of Phosphorothioate 2′-O-(2-Methoxyethyl)-Modified Antisense Oligonucleotides in Animals and Man. In *Antisense Drug Technology: Principles, Strategies, and Applications*, Second Edition; 2007. 10.1201/9780849387951.ch11.

(16) Gagliardi, M.; Ashizawa, A. T. The Challenges and Strategies of Antisense Oligonucleotide Drug Delivery. Biomedicines. 2021. 10.3390/biomedicines9040433.

(17) Sharma, S.; Borski, C.; Hanson, J.; Garcia, M. A.; Link, C. D.; Hoeffer, C.; Chatterjee, A.; Nagpal, P. Identifying Optimal Neuroinflammation Treatment Using Nanoligomer^TM^ Discovery Engine. ACS Chem. Neurosci. 2022, 13 (23), 3247–3256. 10.1021/acschemneuro.2c00365.

(18) Courtney, C. M.; Sharma, S.; Fallgren, C. M.; Weil, M. M.; Chatterjee, A.; Nagpal, P. Reversing Radiation-Induced Immunosuppression Using a New Therapeutic Modality. Life Sci. Sp. Res. 2022, 35, 127–139. 10.1016/j.lssr.2022.05.002.

(19) McCollum, C. R.; Courtney, C. M.; O’Connor, N. J.; Aunins, T. R.; Jordan, T. X.; Rogers, K. L.; Brindley, S.; Brown, J. M.; Nagpal, P.; Chatterjee, A. Safety and Biodistribution of Nanoligomers Targeting the SARS-CoV-2 Genome for the Treatment of COVID-19. ACS Biomater. Sci. Eng. 2023, 9 (3), 1656–1671. 10.1021/acsbiomaterials.2c00669.

(20) McCollum, C. R.; Courtney, C. M.; O’Connor, N. J.; Aunins, T. R.; Ding, Y.; Jordan, T. X.; Rogers, K. L.; Brindley, S.; Brown, J. M.; Nagpal, P.; Chatterjee, A.; O’Connor, N. J.; Aunins, T. R.; Ding, Y.; Jordan, T. X.; Rogers, K. L.; Brindley, S.; Brown, J. M.; Nagpal, P.; Chatterjee, A. NanoligomersTM Targeting Human MiRNA for Treatment of Severe COVID-19 Are Safe and Nontoxic in Mice. ACS Biomater. Sci. Eng. 2022, 8 (7), 3087– 3106. 10.1021/acsbiomaterials.2c00510.

(21) McDonald, J. T.; Enguita, F. J.; Taylor, D.; Griffin, R. J.; Priebe, W.; Emmett, M. R.; Sajadi, M. M.; Harris, A. D.; Clement, J.; Dybas, J. M.; Aykin-Burns, N.; Guarnieri, J. W.; Singh, L. N.; Grabham, P.; Baylin, S. B.; Yousey, A.; Pearson, A. N.; Corry, P. M.; Saravia-Butler, A.; Aunins, T. R.; Sharma, S.; Nagpal, P.; Meydan, C.; Foox, J.; Mozsary, C.; Cerqueira, B.; Zaksas, V.; Singh, U.; Wurtele, E. S.; Costes, S. V.; Davanzo, G. G.; Galeano, D.; Paccanaro, A.; Meinig, S. L.; Hagan, R. S.; Bowman, N. M.; Wallet, S. M.; Maile, R.; Wolfgang, M. C.; Mock, J. R.; Torres-Castillo, J. L.; Love, M. K.; Lovell, W.; Rice, C.; Mitchem, O.; Burgess, D.; Suggs, J.; Jacobs, J.; Altinok, S.; Sapoval, N.; Treangen, T. J.; Moraes-Vieira, P. M.; Vanderburg, C.; Wallace, D. C.; Schisler, J. C.; Mason, C. E.; Chatterjee, A.; Meller, R.; Beheshti, A.; Bowen, R. A.; Griffin, R. J.; Priebe, W.; Emmett, M. R.; McGrath, M.; Sajadi, M. M.; Harris, A. D.; Clement, J.; Dybas, J. M.; Aykin-Burns, N.; Guarnieri, J. W.; Singh, L. N.; Grabham, P.; Baylin, S. B.; Yousey, A.; Pearson, A. N.; Corry, P. M.; Saravia-Butler, A.; Aunins, T. R.; Nagpal, P.; Meydan, C.; Foox, J.; Mozsary, C.; Cerqueira, B.; Zaksas, V.; Singh, U.; Wurtele, E. S.; Costes, S. V.; Galeano, D.; Paccanaro, A.; Meinig, S. L.; Hagan, R. S.; Bowman, N. M.; Wolfgang, M. C.; Altinok, S.; Sapoval, N.; Treangen, T. J.; Frieman, M.; Vanderburg, C.; Wallace, D. C.; Schisler, J. C.; Mason, C. E.; Chatterjee, A.; Meller, R.; Beheshti, A. Role of MiR-2392 in Driving SARS-CoV-2 Infection. Cell Rep. 2021, 37 (3), 109839. 10.1016/j.celrep.2021.109839.

(22) Risen, S. J.; Boland, S.; Sharma, S.; Weismann, G.; Shirley, P.; Latham, A.; Hay, A. J. D.; Gilberto, V.; Hines, A. D.; Brindley, S.; Brown, J. M.; McGrath, S.; Chatterjee, A.; Nagpal, P.; Moreno, J. A. Targeting Neuroinflammation by Pharmacologic Down-Regulation of Inflammatory Pathways Is Neuroprotective in Protein Misfolding Disorders. ACS Chem. Neurosci. 2024. 10.1101/2022.09.26.509513.

(23) Sharma, S.; Risen, S.; Gilberto, V.; Boland, S.; Chatterjee, A.; Moreno, J.; Nagpal, P. Targeted-Neuroinflammation Mitigation Using Inflammasome-Inhibiting Nanoligomers Is Therapeutic in Experimental Autoimmune Encephalomyelitis (EAE) Mouse Model. bioRxiv 2024. 10.1101/2024.01.11.575262.

(24) Gagliardi, C.; Bunnell, B. A. Large Animal Models of Neurological Disorders for Gene Therapy. ILAR Journal. 2009. 10.1093/ilar.50.2.128.

(25) Sharma, S.; Gilberto, V. S.; Levens, C. L.; Chatterjee, A.; Kuhn, K. A.; Nagpal, P. Microbiome- and Host Inflammasome-Targeting Inhibitor Nanoliogmers Are Therapeutic in Murine Colitis Model. bioRxiv 2024. 10.1101/2024.02.20.581256.

(26) Wahl, D.; Risen, S. J.; Osburn, S. C.; Emge, T.; Sharma, S.; Gilberto, V. S.; Chatterjee, A.; Nagpal, P.; Moreno, J. A.; LaRocca, T. J. Nanoligomers Targeting NF-ΚB and NLRP3 Reduce Neuroinflammation and Improve Cognitive Function with Aging and Tauopathy. bioRxiv 2024. 10.1101/2024.02.03.578493.

(27) Kilså Jensen, K.; Ørum, H.; Nielsen, P. E.; Nordén, B. Kinetics for Hybridization of Peptide Nucleic Acids (PNA) with DNA and RNA Studied with the BIAcore Technique. Biochemistry 1997. 10.1021/bi9627525.

(28) Fransson, J.; Espander-Jansson, A. Local Tolerance of Subcutaneous Injections. J. Pharm. Pharmacol. 1996, 48 (10). 10.1111/j.2042-7158.1996.tb05892.x.

(29) ReaganLJShaw, S.; Nihal, M.; Ahmad, N. Dose Translation from Animal to Human Studies Revisited. FASEB J. 2008, 22 (3). 10.1096/fj.07-9574lsf.

(30) Zhang, Y.; Huo, M.; Zhou, J.; Xie, S. PKSolver: An Add-in Program for Pharmacokinetic and Pharmacodynamic Data Analysis in Microsoft Excel. Comput. Methods Programs Biomed. 2010, 99 (3). 10.1016/j.cmpb.2010.01.007.

(31) Blanchard, O. L.; Smoliga, J. M. Translating Dosages from Animal Models to Human Clinical Trials-Revisiting Body Surface Area Scaling. FASEB Journal. 2015. 10.1096/fj.14-269043.

(32) Larsen, H. J.; Bentin, T.; Nielsen, P. E. Antisense Properties of Peptide Nucleic Acid. Biochim. Biophys. Acta - Gene Struct. Expr. 1999, 1489 (1), 159–166. 10.1016/S0167-4781(99)00145-1.

