## Supporting Information for "Large- and Small-Animal Studies of Safety, Pharmacokinetics (PK), and Biodistribution of Inflammasome-Targeting Nanoligomer in the Brain and Other Target Organs"

**Sample ID	Below	p-value**

**Threshold**

| G-CSF (CSF-3) | No | 0.570115 |
| --- | --- | --- |
| IL-10 | No | 0.876126 |
| IL-3 | No | 0.795409 |
| LIF | No | 0.419316 |
| IL-1 beta | No | 0.353403 |
| IL-2 | No | 0.238137 |
| M-CSF | No | 0.899768 |
| IP-10 (CXCL10) | No | 0.375312 |
| IL-4 | No | 0.576022 |
| IL-5 | No | 0.932894 |
| IL-6 | No | 0.895916 |
| IL-22 | No | 0.227429 |
| IL-9 | No | 0.230025 |
| IL-13 | No | 0.753696 |
| IL-27 | No | 0.560482 |
| IL-23 | No | 0.396848 |
| IFN gamma | No | 0.706989 |
| IL-12p70 | No | 0.252759 |
| GM-CSF | No | 0.440844 |
| GRO alpha (CXCL1) | No | 0.384355 |
| RANTES (CCL5) | No | 0.355962 |
| TNF alpha | No | 0.897307 |
| MIP-1 alpha (CCL3) | No | 0.799229 |
| MCP-3 (CCL7) | No | 0.713972 |
| MCP-1 (CCL2) | No | 0.680433 |
| IL-17 A (CTLA-8) | No | 0.531265 |
| IL-15 | No | 0.519355 |
| MIP-2 alpha (CXCL2) | No | 0.477553 |
| IL-1 alpha | No | 0.540948 |
| ENA-78 (CXCL5) | No | 0.820599 |
| Eotaxin (CCL11) | No | 0.409270 |
| IL-28 | No | 0.973117 |
| IL-18 | No | 0.724024 |
| MIP-1 beta (CCL4) | No | 0.788846 |
| IL-31 | No | 0.413313 |

**Fig. S1.** p-values for statistical comparison between NI112 treated and sham-treated animals assessed using blood serum with 36-plex cytokine and chemokine panel, using unpaired t-test for each cytokine.

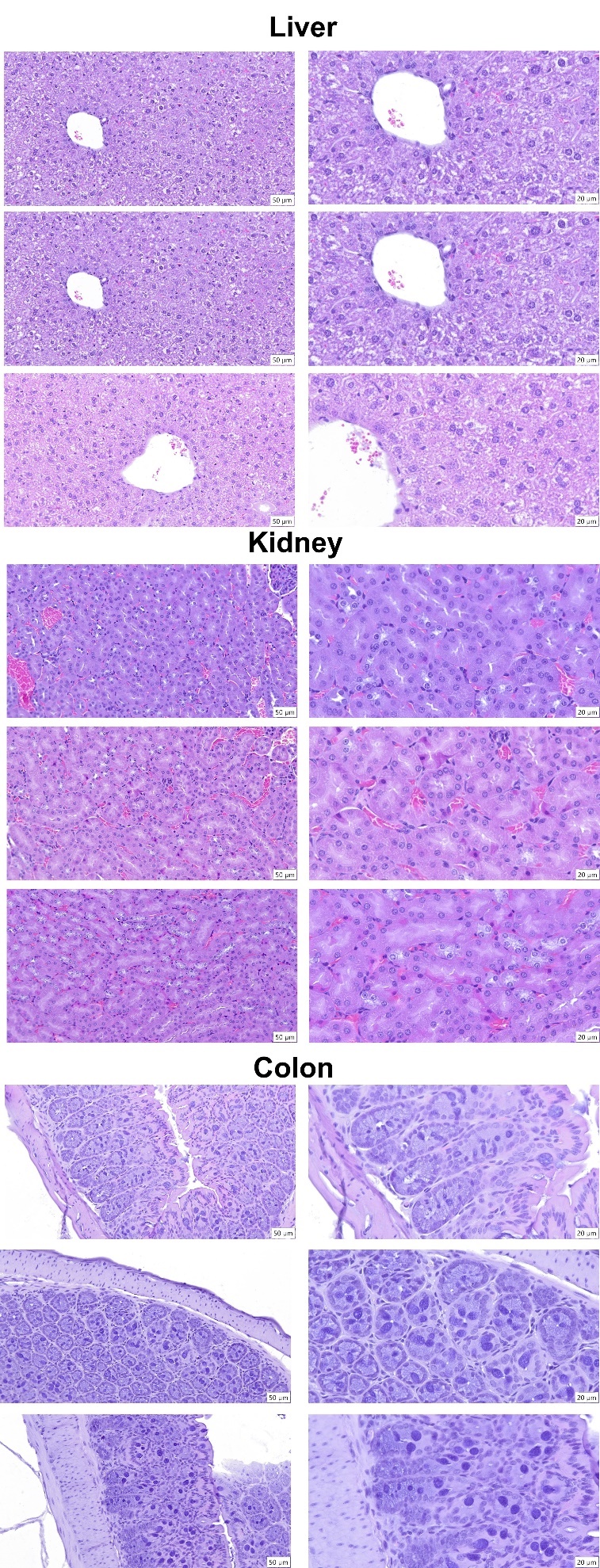
**Fig. S2.** Additional H&E-stained histology of liver, kidney, and colon samples from mice after long-term (12 weeks) treatment. Histologic findings in each tissue were diagnosed and graded for severity on a 0-5 scale based on inflammation and hemorrhage. 0=absent, 1=minimal (10% of tissue affected), 2=mild (10-25% of tissue affected), 3=moderate (26-50% of tissue affected), 4=marked (51-75% of tissue affected), 5=severe (75% of tissue affected). The middle kidney histology showed mild hemorrhage and was scored at 1. The remaining samples (using further images) were scored as zero, as shown in **Fig. 1B**. There were no significant histologic findings in the liver, kidney, or colons of mice treated with NI112. n=4 animal/group.

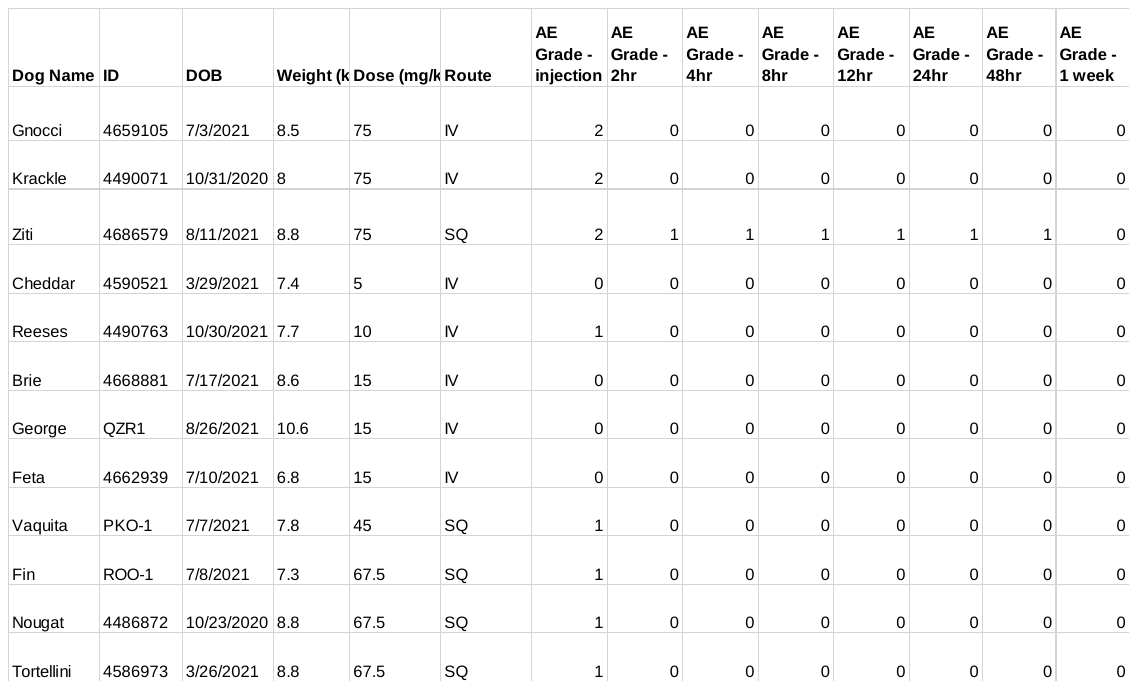

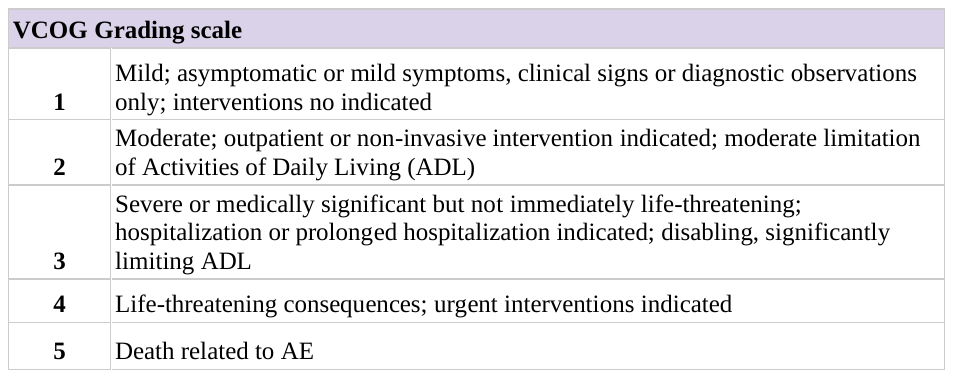
**Fig. S3.** (Top) Clinical evaluation data summary for 12 dogs enrolled in the safety-toxicity study, using veterinary cooperative oncology group-common terminology criteria for adverse events (VOG-CTCAE). Adverse events (AE) were documented following API administration and used to determine MTD. The SQ-dosed dogs (IDs PKO-1, ROO-1, 4486872, and 4586973) scored 1 as AE registered mild scratching at the injection site, maximum up to 5-10 minutes. The subjects were fine during the remaining evaluation period (15 minutes) and resumed normal lifestyle. Further, AE evaluations during the day did not observe any changes from normal (scored AE=0 in subsequent examinations). For the above MTD evaluations, dog ID 4659105 showed extreme nausea, loose stool, and vomiting post-injection, but no AE after subsequent evaluations (scored zero). 10 minutes post-injection, subject ID 4490071 had a brief seizure/convulsion lasting 10 seconds and urinated, but then resumed normal function. These two AEs indicated 75 mg/kg dose as above MTD. For SQ, dog ID 4686579 had pain immediately after injection, and the pain reaction lasted 2 minutes, with the injection site painful to touch 5 minutes post-injection. This AE was later traced back to the high pH of SQ injection^26^ due to the use of unbuffered saline media to dissolve dried NI112 powder. However, as per IACUC protocol at CSU, the AE resulted in 75mg/kg marked as above MTD. As seen in subsequent CBC/chemistry, and clinical evaluations, no organ damage or pathological events were seen at the highest 75 mg/kg dosages (**Fig. 1C-F**, **S4, S5**).

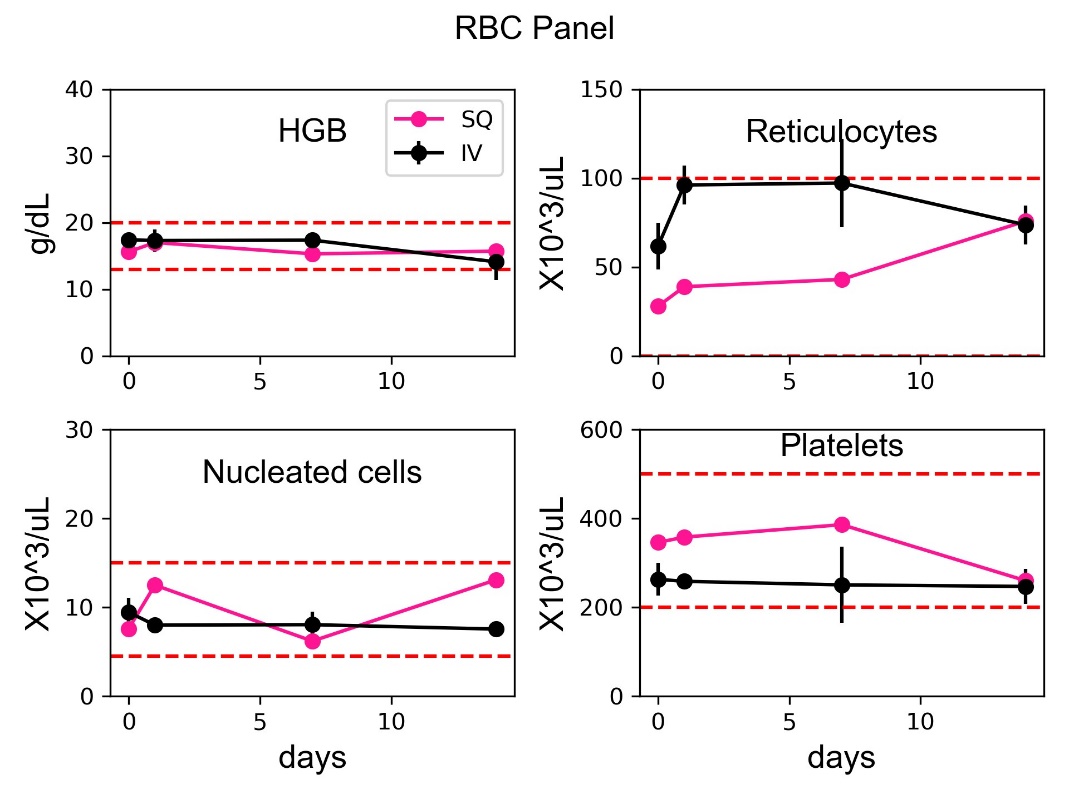

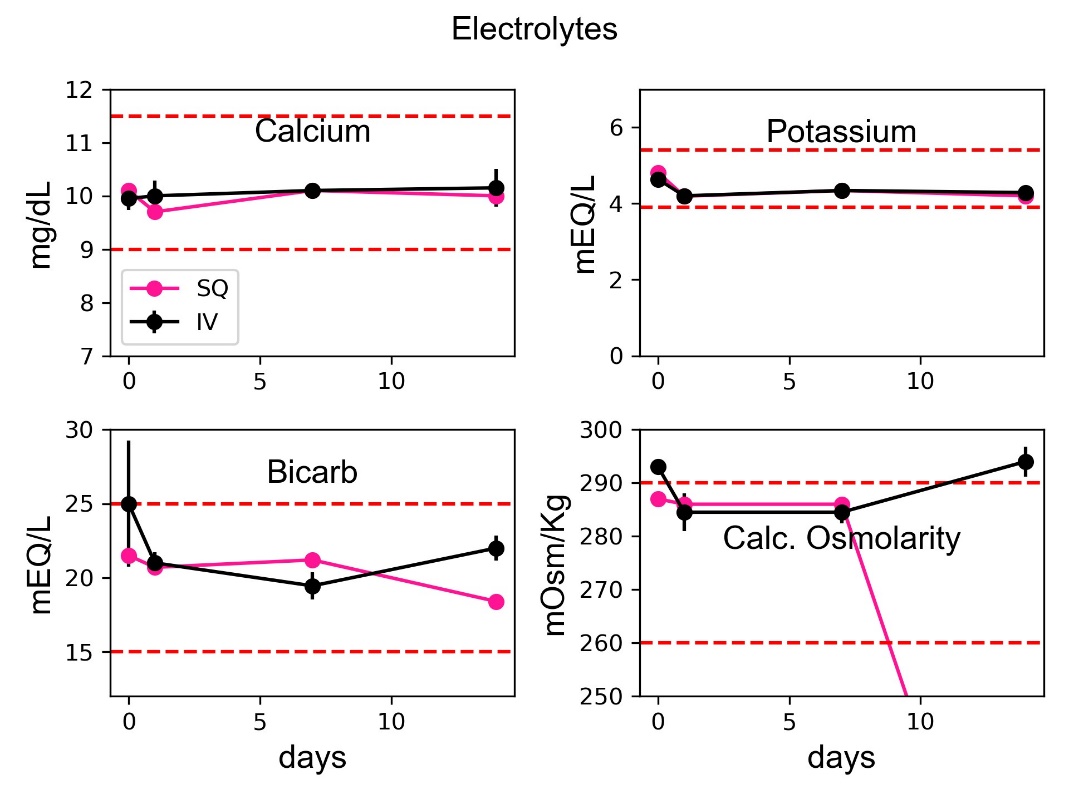

**Fig. S4**. Monitoring health using Red Blood Cells (RBC) panel (Top) and Electrolyte panel (Bottom). RBC panel includes Hemoglobin (HGB), reticulocyte nucleated cells, and platelets, whereas the Electrolyte panel includes calcium, potassium, bicarbonate, and calc. osmolarity, with extensive CBC/chemistry monitoring over 14 days using three clinically translatable routes of drug administration (intravenous IV black, subcutaneous SQ pink), after administration of the highest dose (75 mg/kg dose IV and SQ) well above MTD. The red dashed lines show upper and lower normal limits for each measurement. The curves show mean values, and the bars show standard deviation (Mean ± SD).

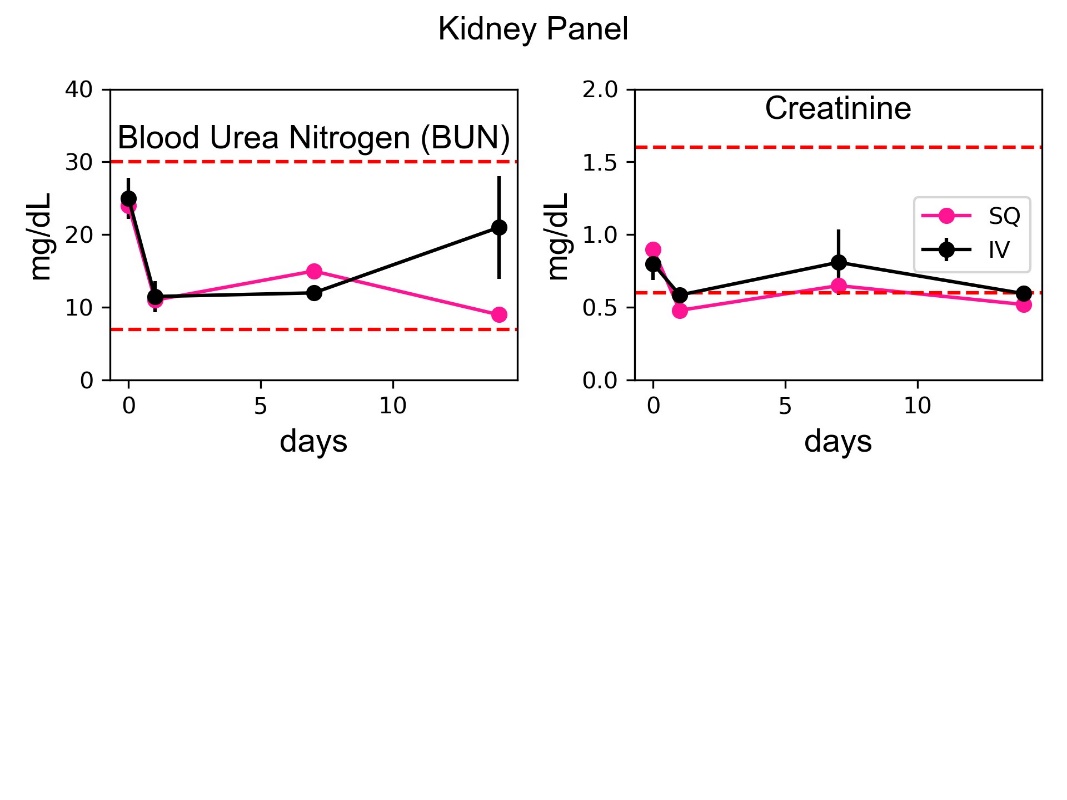

**Fig. S5.** Monitoring kidney health using blood urea nitrogen, and creatinine, as seen using extensive CBC/chemistry monitoring over 14 days using three clinically translatable routes of drug administration (intravenous IV black, subcutaneous SQ pink), after administration of the highest dose (75 mg/kg dose IV and SQ) well above MTD. The red dashed lines show upper and lower normal limits for each measurement. The curves show mean values, and the bars show standard deviation (Mean ± SD).

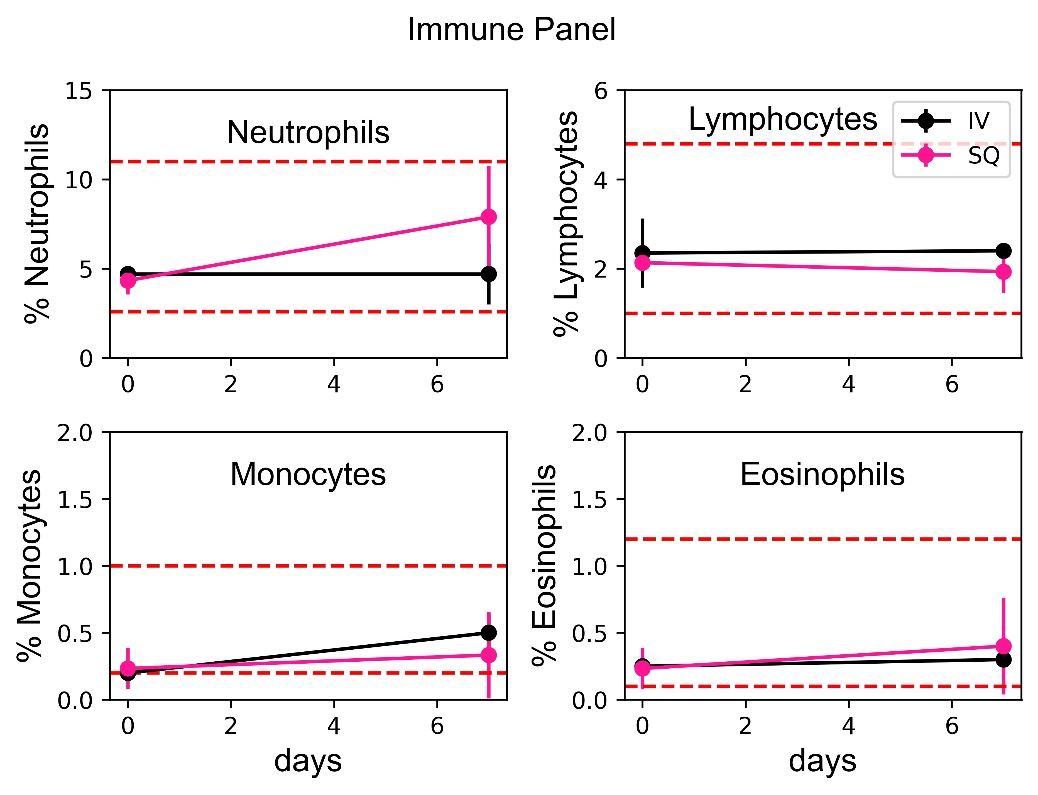

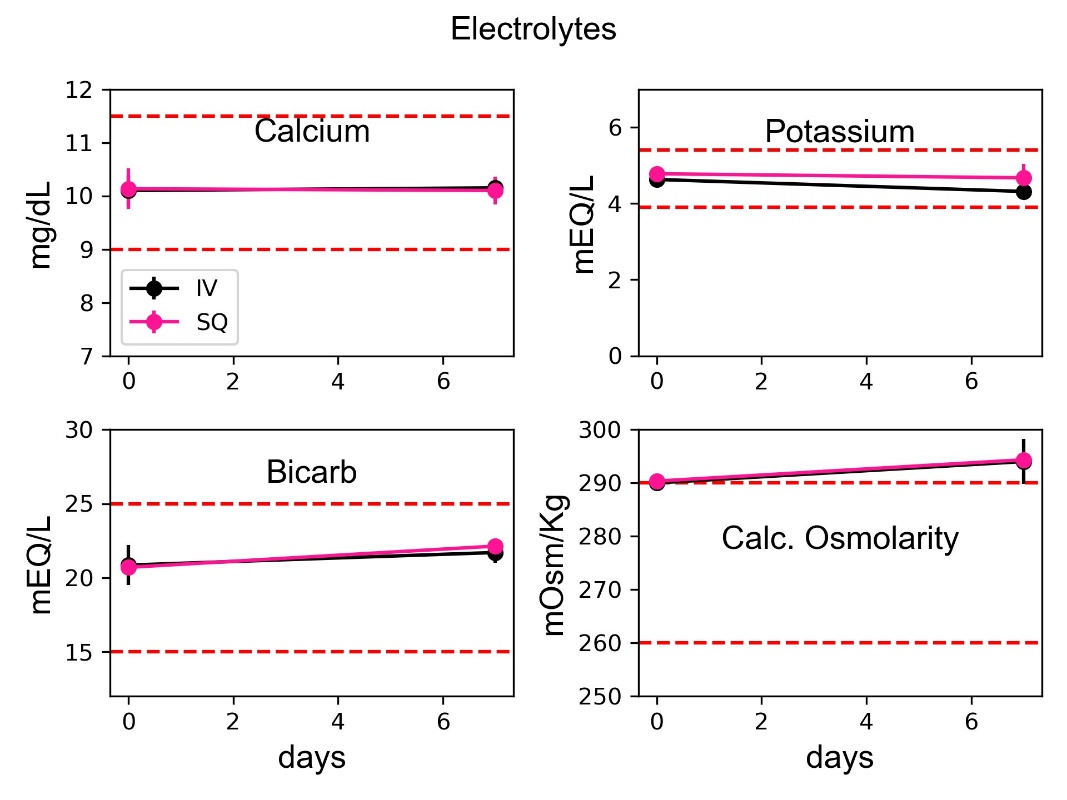

**Fig. S6.** Lack of any immunogenic response, or imbalance in electrolyte concentrations can be seen using extensive CBC/chemistry monitoring over 7 days after administration of 15 mg/kg and 67.5 mg/kg MTD dosing using IV and SQ routes, respectively. The red dashed lines show upper and lower normal limits for each measurement. Using monitoring of different immune components and total blood count, all immune biomarkers were found to be normal. The curves show mean values, and the bars show standard deviation (Mean ± SD).

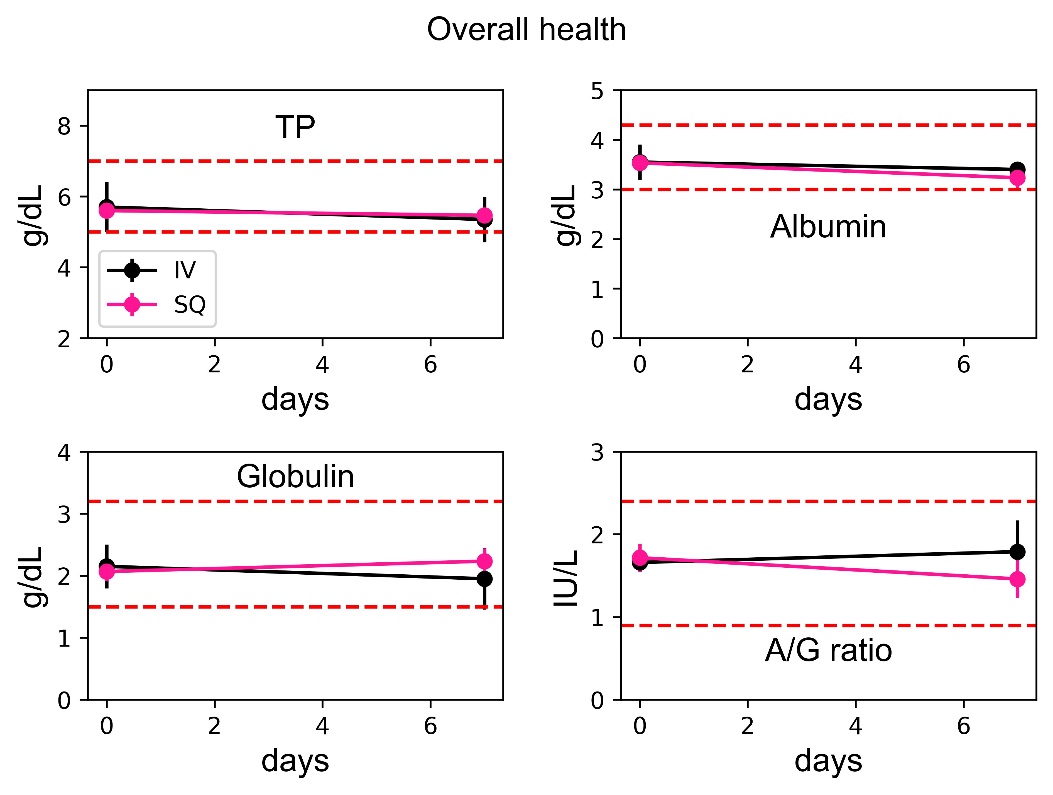

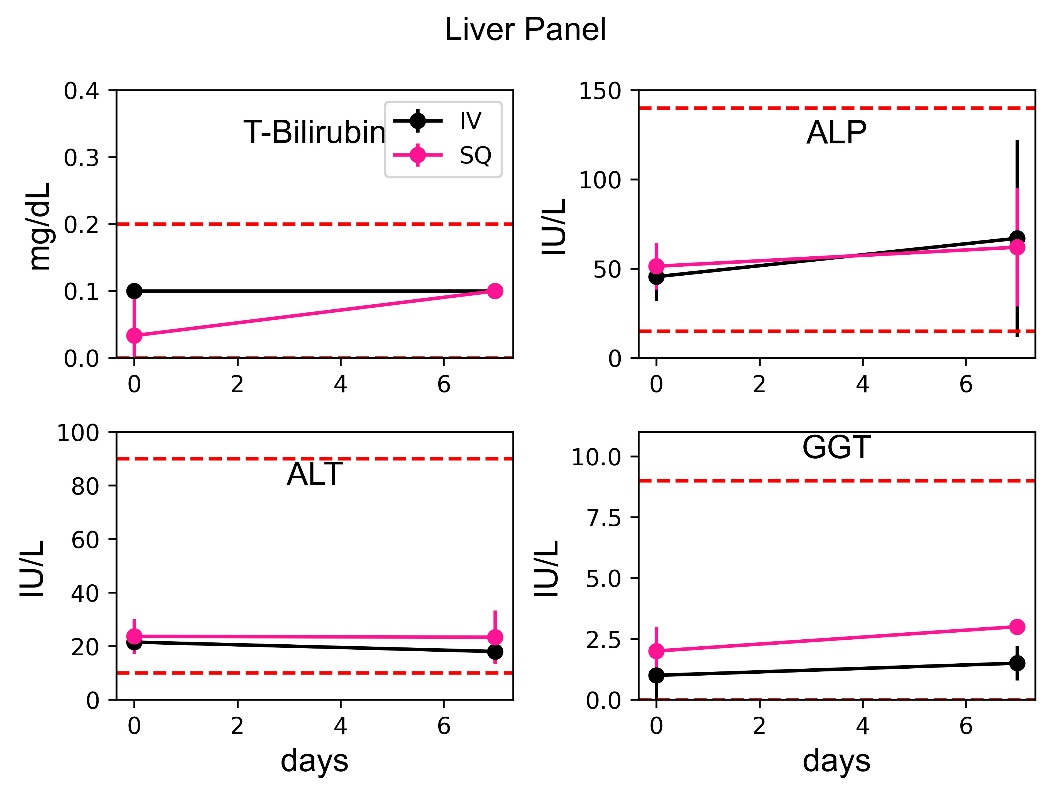

**Fig. S7.** Lack of any adverse impact on the overall health (using total protein (TP), albumin, globulin, Albumin, and A/G ratio), or liver health (using total bilirubin, ALP), ALT, and GGT enzymes), can be seen using extensive CBC/chemistry monitoring over 7 days after administration of 15 mg/kg and 67.5 mg/kg MTD dosing using IV and SQ routes, respectively. The red dashed lines show upper and lower normal limits for each measurement. The curves show mean values, and the bars show standard deviation (Mean ± SD).

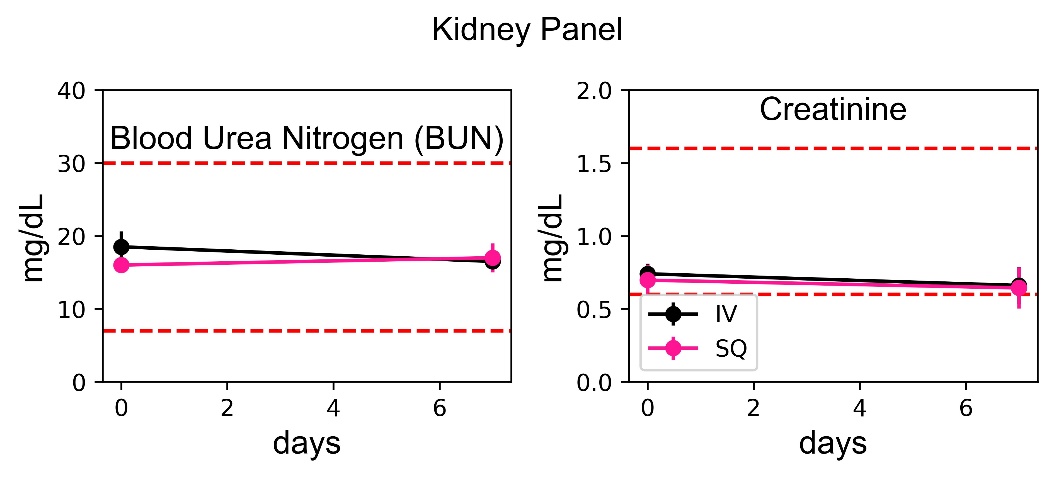

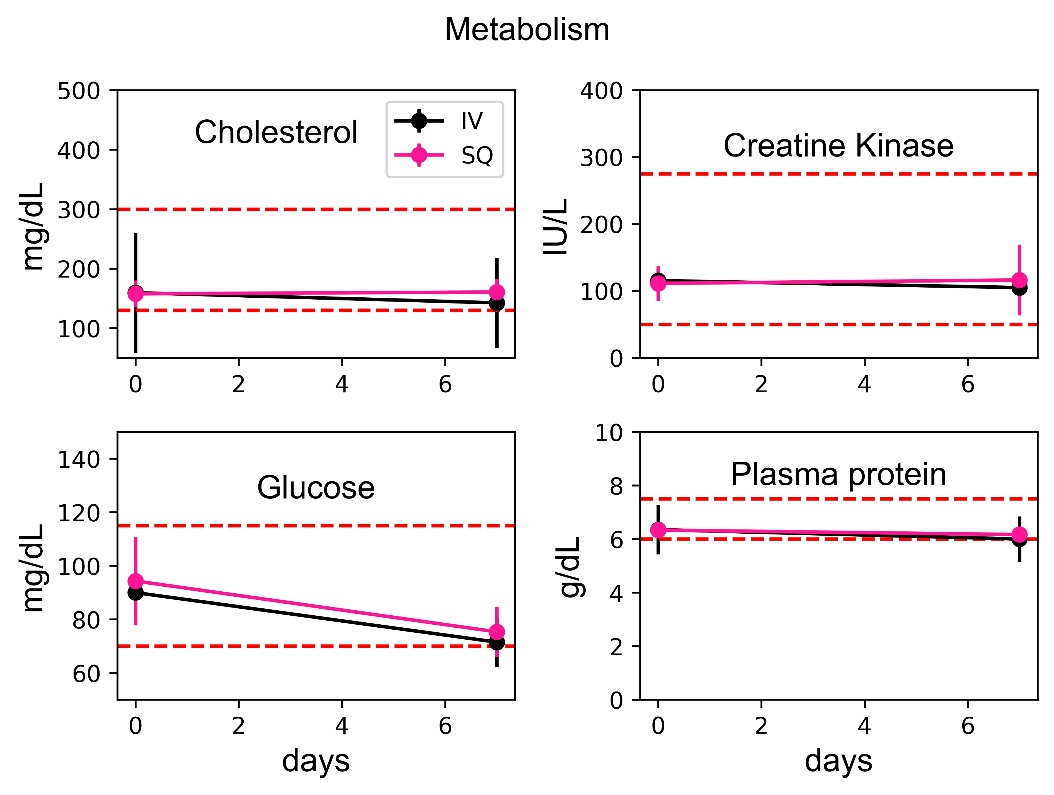

**Fig. S8.** Lack of any adverse changes in kidney health (using blood urea nitrogen, and creatinine), and metabolism and cardiac health (using cholesterol, creatinine kinase, glucose, and plasma protein), can be seen using extensive CBC/chemistry monitoring over 7 days after administration of 15 mg/kg and 67.5 mg/kg MTD dosing using IV and SQ routes, respectively. The red dashed lines show upper and lower normal limits for each measurement. The curves show mean values, and the bars show standard deviation (Mean ± SD).

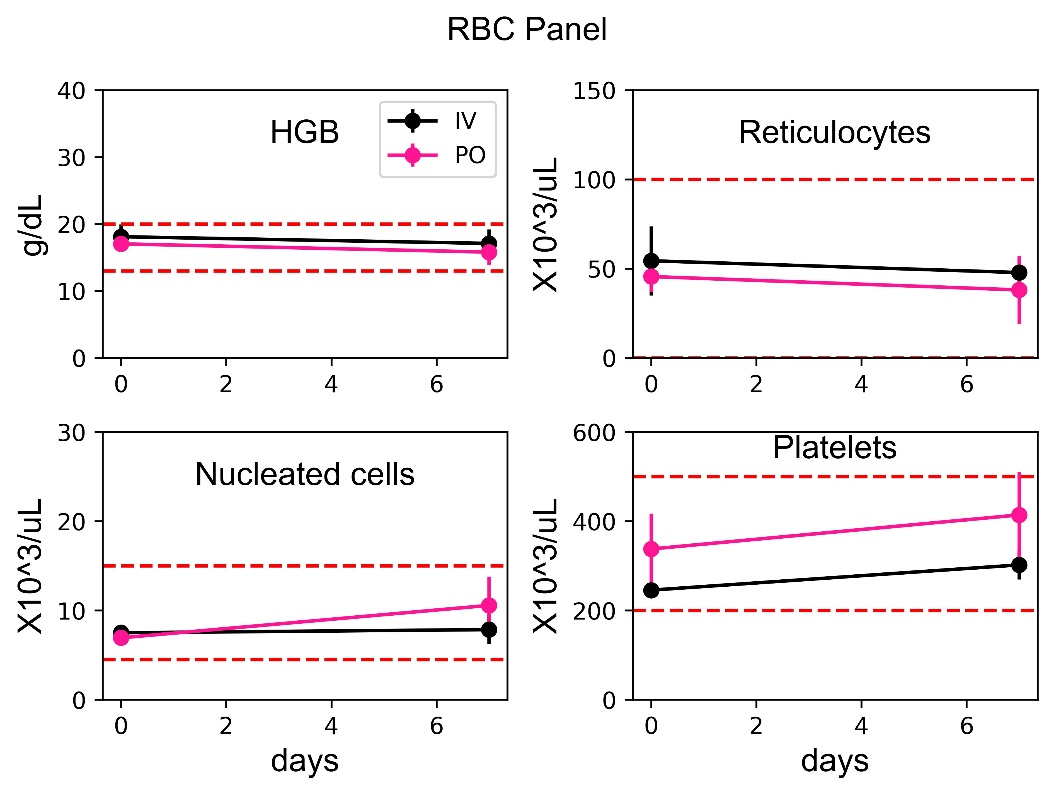

**Fig. S9.** Lack of any adverse changes in the RBC panel (Hemoglobin (HGB), reticulates, nucleated cells, and platelets) can be seen using extensive CBC/chemistry monitoring over 7 days after administration of 15 mg/kg and 67.5 mg/kg MTD dosing using IV and SQ routes, respectively. The red dashed lines show upper and lower normal limits for each measurement. The curves show mean values, and the bars show standard deviation (Mean ± SD).

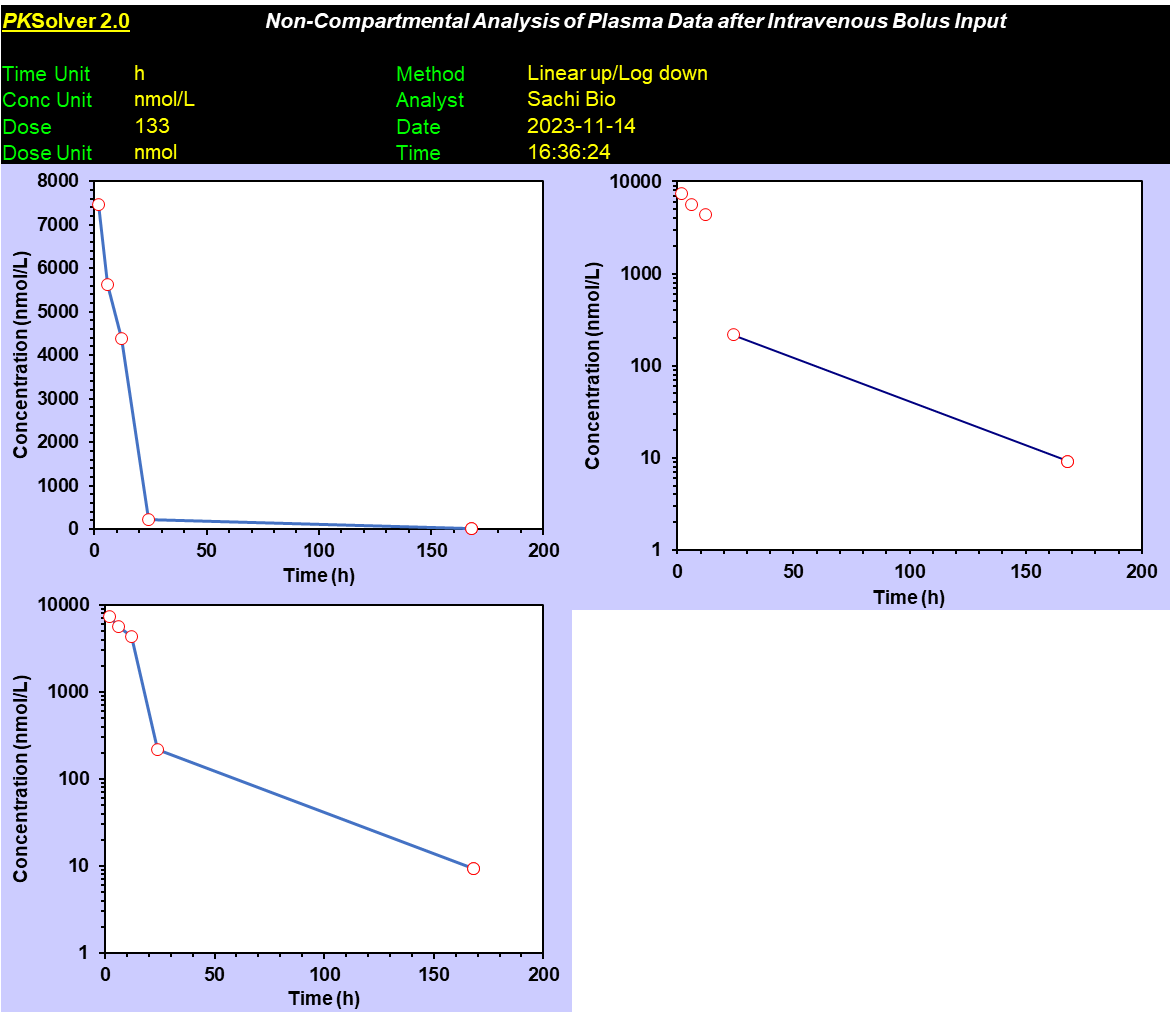

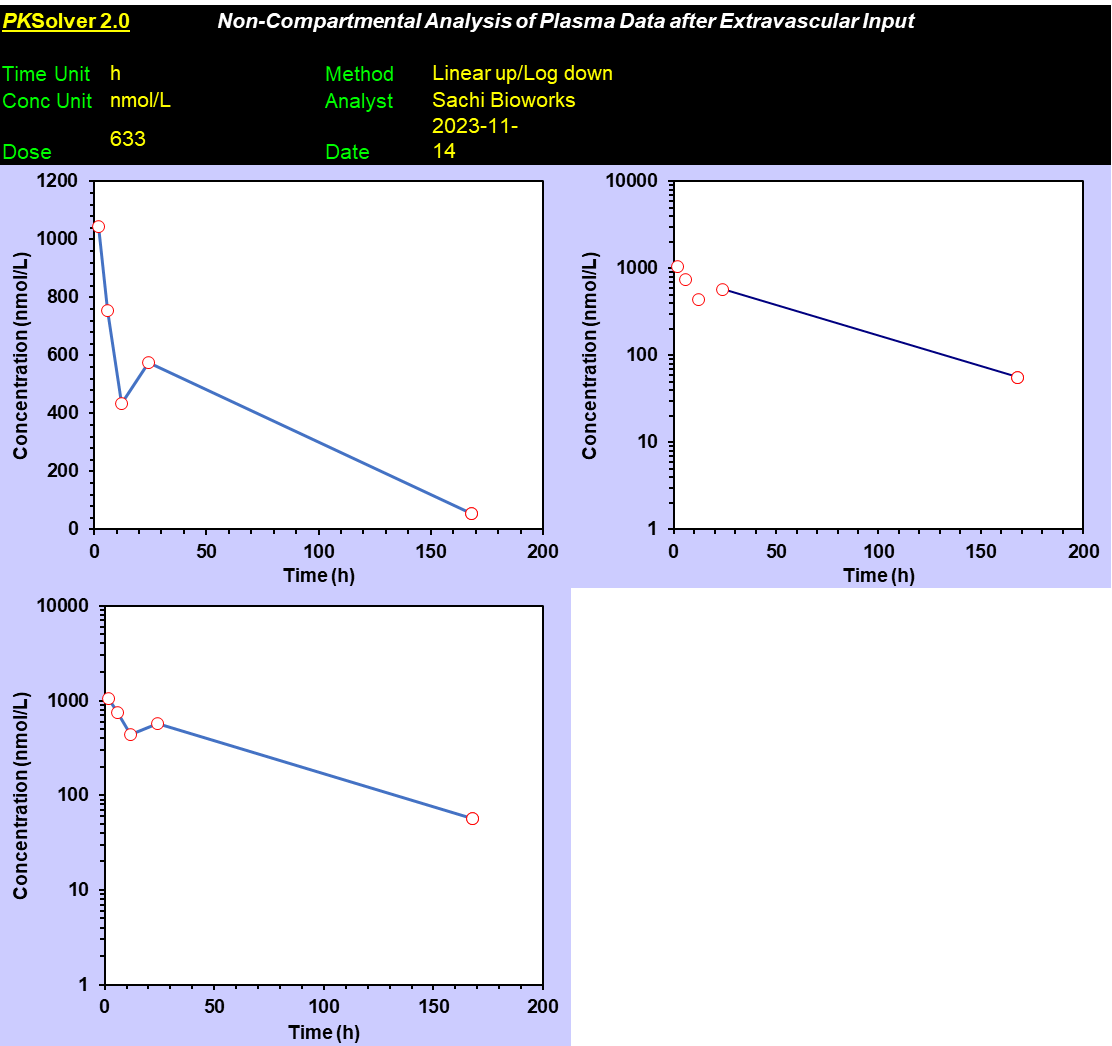

**Fig. S10**. **Pharmacokinetic modeling** using PK Solver^28^ for mice using the IV route (Top) and SQ route (bottom) of administration. n=4 mice per group, Median ± SE was used for analysis (**Fig.2**).

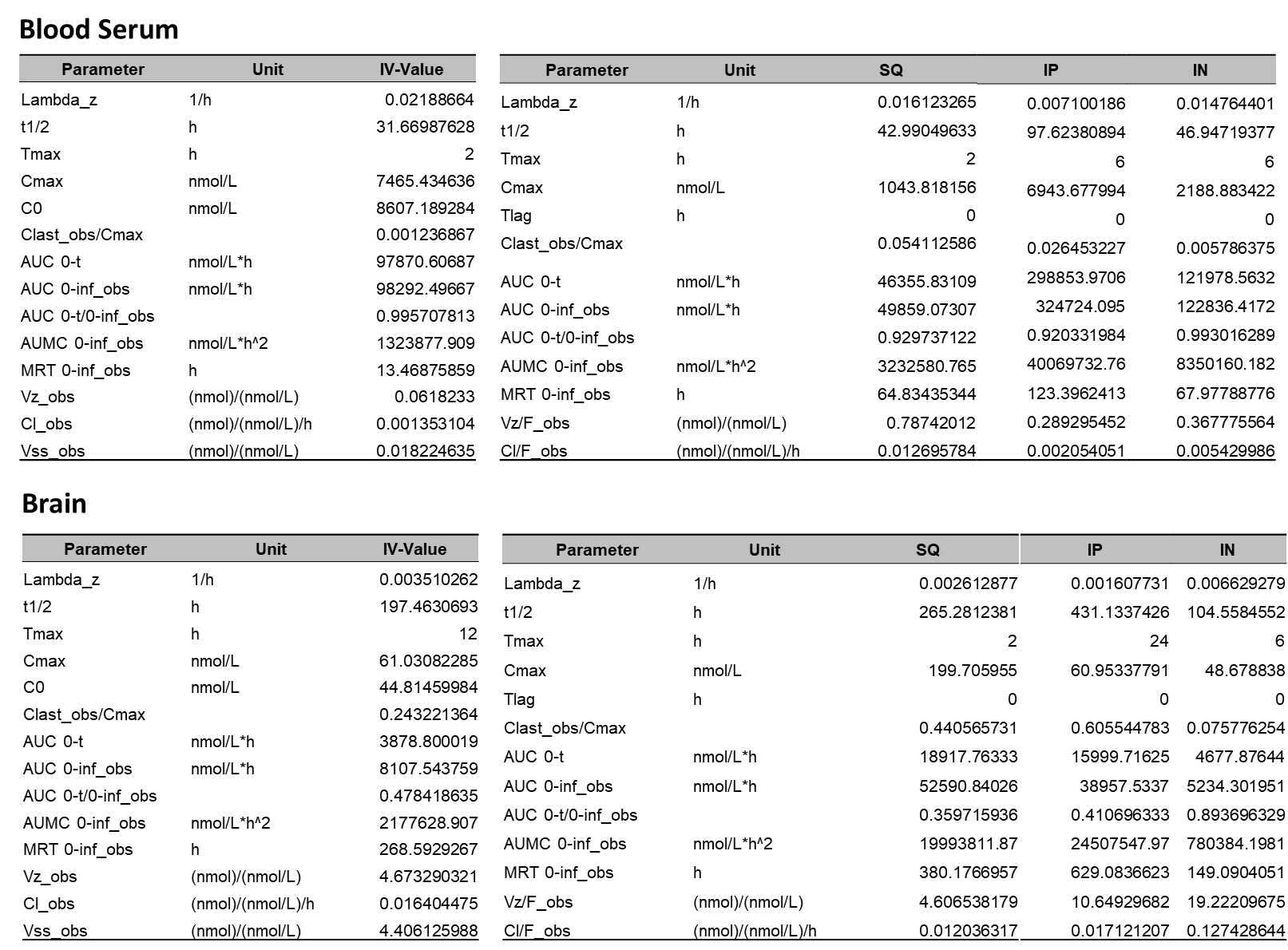

**Fig. S11**. **Pharmacokinetic modeling parameters** obtained using PK Solver^28^ for mice using IV (n=4/group), SQ (n=4/group), IP (n=6/group), and IN (n=6/group) in (Top) blood serum; and (Bottom) brain tissue. Median ± SE used for average and SE analysis shown in **Figs.2, 3, 4**.

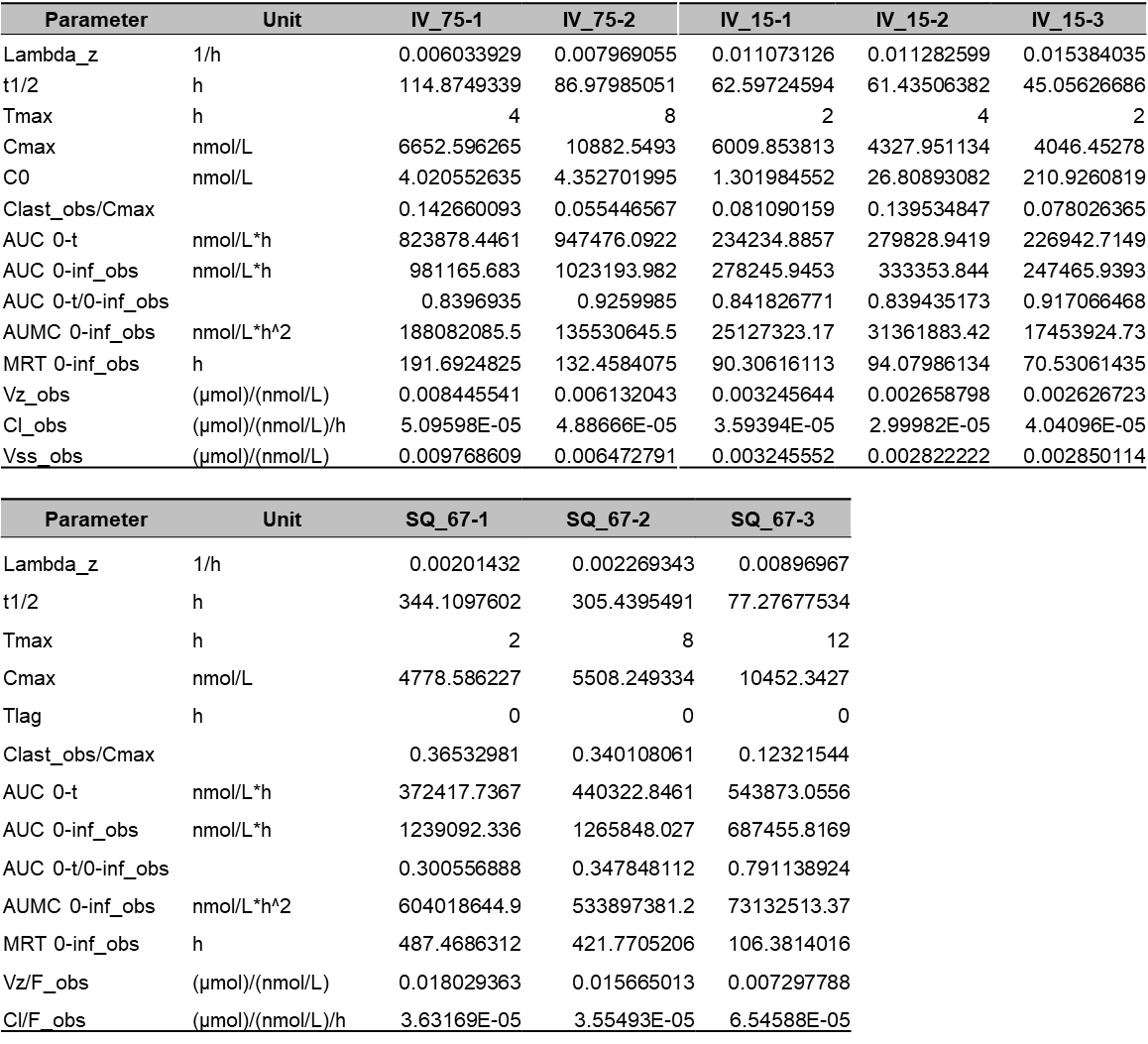

**Fig. S12**. **Pharmacokinetic modeling parameters** obtained using PK Solver^28^ for dogs using IV MTD (15 mg/kg), IV acute dose (75 mg/kg, 5x MTD), and SQ (67.5 mg/kg).
